# A Direct DNA Extraction Workflow for Metabarcoding Fungal Bioaerosols from Adhesive Samplers

**DOI:** 10.64898/2026.01.12.698475

**Authors:** Jiwon Jung, Rodrigo Leitao, Alice L. F. Day, Huw Davis, Nichola Hawkins, Kostya Kanyuka, Matthew C. Fisher

## Abstract

Monitoring fungal bioaerosols is essential for understanding their impacts on human health and crop productivity. However, current monitoring approaches rely on culture-dependent techniques, which underestimate the diversity of fungal exposures. Adhesive-coated films can effectively capture airborne particles but have been regarded as unsuitable for DNA downstream analysis, owing to the lack of standardised DNA extraction methods for the adhesive capture surface. Here, we present an optimised DNA extraction protocol for adhesive film, which we developed and validated to enable direct and culture-independent ITS2-based metabarcoding of fungal bioaerosols collected on a 96 well plate sealing film (‘Sticky sampler’). This validated approach was applied to field samples collected from four agricultural farms and from a domestic garden to evaluate seasonal and spatial changes in fungal bioaerosols. Our analysis uncovered pronounced site-specific contrasts linked to geography and land management, as well as clear seasonal shifts in garden fungal communities. This study demonstrates that adhesive samplers, combined with optimised DNA extraction, provide a practical and inexpensive tool for fungal bioaerosol research by resolving ecologically meaningful spatiotemporal dynamics of airborne fungi. This protocol opens new opportunities for highly scalable surveillance of environmental bioaerosols.

## 1. Introduction

Fungal bioaerosols represent a major component of airborne biological particles and are primarily released through the dispersal of spores, the dominant form of airborne fungi [1]. Fungal spores are ubiquitous in both indoor and outdoor environments [2]. The composition and concentration of fungal bioaerosols are shaped by environmental factors such as temperature, humidity, and wind patterns. Consequently, fungal bioaerosols display distinct spatial and seasonal distribution patterns [3, 4]. Fungal spores, which range in size from less than 1 µm to about 20 µm, are small enough to be easily transported far from their sources and enter the human body via respiratory and digestive tracts, or through the skin [5–8].

Exposure to fungal bioaerosols has been associated with multiple adverse outcomes, including both environmental and clinical settings. In agriculture, fungal bioaerosols contribute to crop yield losses, while in humans they are responsible for causing infectious diseases, respiratory infections, and skin allergies [9–12]. The death of two-year-old Awaab Ishak in 2020, caused by prolonged exposure to indoor mould, strongly highlighted the severe health impacts of fungal bioaerosols and raised public awareness of indoor air quality issues [13]. Given the adverse effects of fungal spores in both agriculture and clinical environments, the development of reliable detection and characterisation methods for fungal bioaerosols is crucial for accurate bioaerosol monitoring and health risk assessments.

Effective air sampling forms the foundation for reliable monitoring and subsequent characterisation of fungal bioaerosols. Despite its importance, establishing a robust sampling protocol remains challenging. This difficulty arises because the choice of sampling method largely depends on particle size, yet fungal bioaerosols span a wide size range [12]. Although many types of samplers exist, analytical results can vary depending on the device used, owing to differences in collection efficiency and compatibility with downstream analyses [14]. Since no single air sampling device performs optimally across different particle sizes, analytical techniques are typically selected according to the sampler employed [14]. A further limitation is that culture-based sampling, although still widely used in bioaerosol studies, captures only a small fraction of the fungal community because only an estimated 1–10% of bacteria and fungi grow under standard laboratory conditions [1, 15].

For these reasons, glue-based samplers offer an attractive alternative to traditional culture-based techniques as they are able to capture a broad spectrum of airborne particles, operate without the need for power, and are cost effective, making them suitable for large-scale monitoring. However, the broader application of glue-based samplers in molecular bioaerosol research has been limited by the absence of a standardised protocol for efficient DNA extraction from the adhesive surface. Previous culture-independent attempts have primarily relied on vaseline-or silicone-coated samplers [16].

The ‘Sticky Samplers’ used in this study are sterile adhesive films coated with glue on one side, originally designed for sealing 96-well PCR plates. These samplers were repurposed here as passive bioaerosol collectors, which offer several advantages. They are manufactured under sterile conditions, minimising external contamination during field deployment. In addition, the composition of the sticky samplers confers high thermal and chemical stability typical of PCR sealing materials. Furthermore, their simplicity eliminates the need for extensive technical training, allowing non-experts to conduct large-scale sampling. This accessibility enables participation in citizen-science studies and supports the collection of larger numbers of samplers, thereby enhancing both spatial and temporal coverage in environmental monitoring. Notably, this type of sampler has also been used in fungal studies, demonstrating its practicality as a collection device, although such applications have so far been limited to culture-dependent workflows [17, 18].

In this study, we have developed an optimised DNA-extraction protocol that enables culture-independent metabarcoding of fungal bioaerosols collected on glue-based adhesive films. Using samples collected from farms and gardens, this approach was shown to capture spatiotemporal variation in airborne fungal communities under real-world conditions. This approach expands the applicability of adhesive samplers for large-scale and community-based environmental monitoring, with broad implications for public health, agriculture, and citizen science.

## 2. Methods and Materials

### 2.1 ‘Sticky Sampler’

The adhesive films used as ‘Sticky Samplers’ were Microseal ‘B’ PCR Plate Sealing Films (Bio-Rad Laboratories, USA; catalogue no. MSB1001B). The adhesive surface was left exposed for four weeks for both indoor and outdoor deployments. For indoor environments, the samplers were placed on tables or shelves, whereas for outdoor settings, they were placed within delta traps and hung from tree branches or other suitable structures to protect the adhesive surface from climatic factors such as rain and wind [18]. Before deployment, each film was stored in its original sterile bag with a protective paper layer covering the adhesive surface. For sampling, the protective layer was removed and kept inside the sterile bag during the 4-week exposure. After exposure, the protective layer was reattached, and each sampler was returned to the same sterile bag and stored at room temperature until DNA extraction.

## Method Optimisation

### 2.2 Mock Test: Find an Efficient DNA Extraction Method for Glue Surface

#### 2.2.1 Preparation of Mock Community

To optimise fungal DNA extraction from the sticky samplers, mock tests were performed by spiking them with a fungal mock community consisting of three species: *Aspergillus fumigatus*, *Talaromyces purpurogenus*, and *Talaromyces stollii*. Each species was separately cultured and counted using a haemocytometer. Spore suspensions were adjusted to 2000 spores/µL, mixed in equal volumes, and serially diluted (10⁵–10⁰ spores per 50 µL). Subsequently, 50 µL of each dilution was applied to the centre of a sticky sampler cut to 3 × 1 cm. Spiked samplers were placed in sterile Petri dishes and left to dry overnight in a safety cabinet under controlled conditions.

#### 2.2.2 Evaluation of Two Extraction Approaches

Two extraction strategies were evaluated to identify the most effective method for recovering fungal DNA from the sticky samplers spiked with mock communities.

I Solvent-based method involved removing the adhesive layer with either acetone or ethanol prior to DNA extraction. For each solvent, two incubation periods (20 min and 2 h) were evaluated by immersing the spiked samples in 2 mL of solvent within microcentrifuge tubes and incubating with gentle agitation on a laboratory shaker. After incubation, the solvent was allowed to evaporate under a fume hood and the sticky samplers were discarded. This approach was explored based on the hypothesis that partial dissolution of the glue might improve DNA release from the sticky surface.
II Direct extraction method skipped the adhesive removal step, motivated by an abstract that was presented at a conference [19], which used a different type of tape, double-sided black carbon tape (Carbon Conductive Tape, 25 mm W × 5 m L; Product No. 16084-8, Ted Pella Inc., USA), for sample collection. This approach aimed to assess whether efficient DNA recovery could be achieved without pretreatment for glue removal, thereby simplifying the workflow and reducing sample loss.

Apart from the initial glue removal step in the Solvent-based approach, all subsequent DNA extraction procedures were identical for both methods and are described below.

#### 2.2.3 Optimised DNA Extraction Protocol

The following protocol was used for all subsequent analyses: 800 µL of C1 buffer and the DNA containing sticky sampler (or the DNA residue remaining on the tube walls in the Solvent-based method) were transferred to the bead-containing tubes provided in the DNeasy PowerSoil Pro Kits (Qiagen, Germany). To minimise surface loss and potential contamination, only the edges of the samplers were folded during transfer, thereby ensuring maximum surface contact with the beads inside the tube. The tubes were homogenised using a FastPrep-24™ 5G bead beating grinder and lysis system (MP Biomedicals, USA) with a modified protocol: three cycles at 4.5 m/sec for 45 s, each separated by 15 s of pause, repeated twice in total. After each set of three cycles, the samples were placed on ice to prevent overheating. The volume of C6 was adjusted to 36 µL for this extraction, while remaining steps followed the manufacturer’s protocols. To ensure data validity and monitor potential contamination, the following controls were included during the extraction process: (1) a kit control (KC), consisting of extraction reagents only; and (2) two blank controls, consisting of unused sticky samplers that were either UV-treated (UV) or non-UV-treated (NoUV).

#### 2.2.4 Validation of Method Efficiency

Fungal DNA was quantified by qPCR targeting the 18S rRNA using the FungiQuant primers and probe set [20] on a QuantStudio 7 Flex Real-Time PCR System (v1.7.2 software, Applied Biosystems, USA; Table 1). qPCR was performed in technical triplicate for all samples, including standards, targets, no-template controls (NTCs), the kit control (KC), and the blank controls (UV and NoUV), to ensure consistency and reproducibility. Absolute quantification was performed using a standard curve generated from a 10-fold serial dilution (10⁰–10⁴ genome copy number) of fungal community genomic DNA (MSA-1010, ATCC, USA). As the standard was DNA-based, results are expressed as target gene copy numbers. Copy numbers in the samples were estimated from Ct values based on the slope and y-intercept of the calibration curve, following standard qPCR methods [21, 22]. Since only 2 µL of the final 36 µL DNA extract was used per qPCR reaction, the measured copy numbers were multiplied by 18 (36/2) to estimate the total genome-equivalent copy number in the entire DNA extract.

**Table 1.**
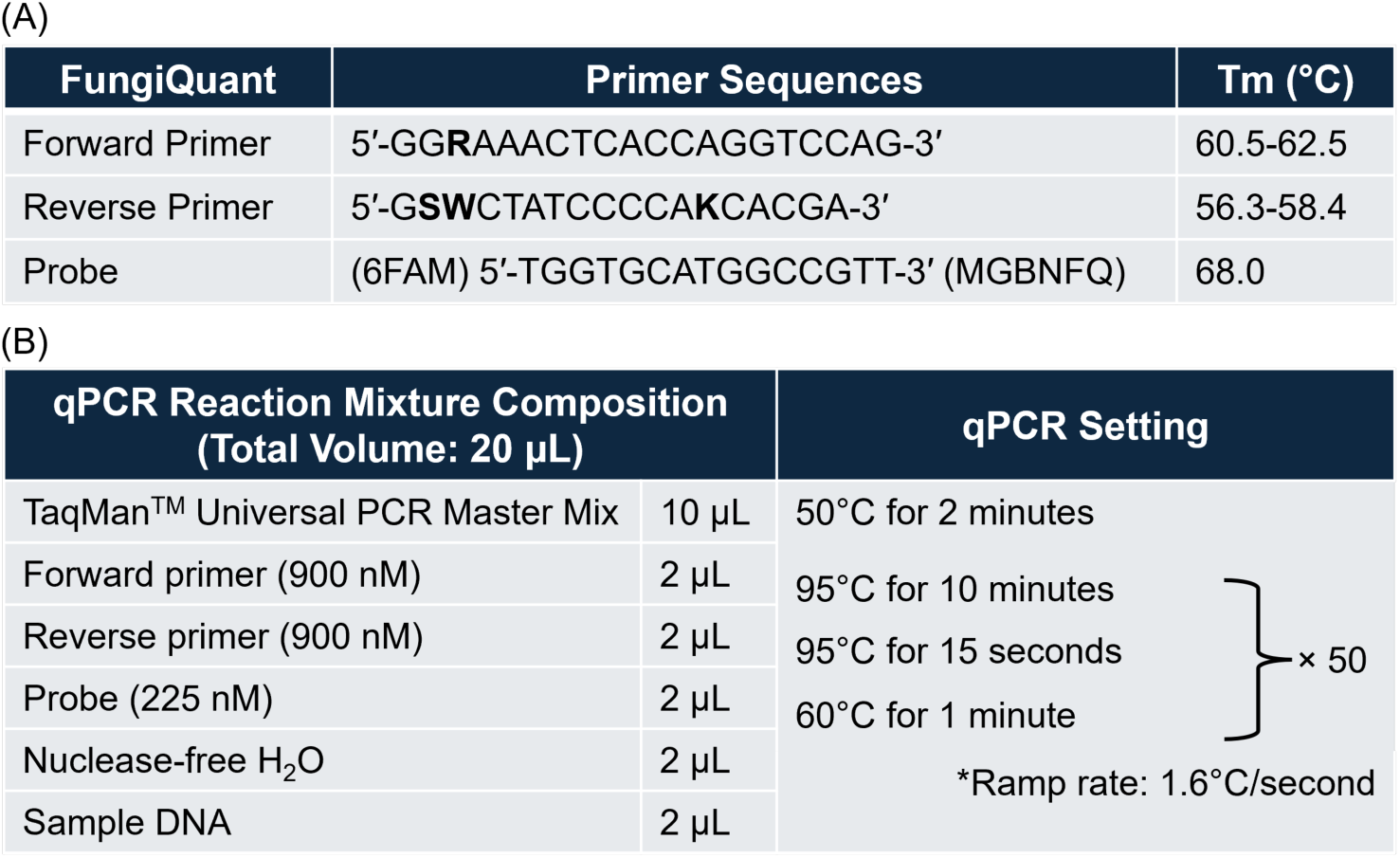
Primers and Probe Sequences and qPCR Conditions Used for the FungiQuant Assay. (A) Primers and probe sequences used in the FungiQuant qPCR assay, including their melting temperatures (Tm). All primers and the probe were synthesised by Thermo Fisher Scientific, USA. (B) Composition of the qPCR reaction mixture and cycling conditions.

The ITS2 region was PCR-amplified using the primer pair ITS7F and ITS4R with DreamTaq PCR Master Mix (Thermo Fisher Scientific, USA). Amplicons were visualised on a 1% agarose gel to confirm DNA amplifiability. The reaction mixture and thermocycling conditions are provided in Table 2.

DNA extractions were performed using the dilution series of the mock fungal mixture (10⁰–10⁵ fungal spores) to evaluate the DNA recovery efficiency of the extraction method. The dilution series was split into two parallel processing groups: (1) Sampling group: aliquots were applied to sticky samplers prior to extraction; and (2) No Sampling group: identical aliquots from the same dilutions were extracted directly without sampler contact to provide a baseline measurement of the original DNA input.

## Method Validation and Sampler Suitability Assessment

### 2.3 Evaluation of Within-Sampler Positional Consistency

Positional consistency was assessed before applying the optimised extraction method to all sticky samplers collected from the environment. This experiment was performed to ensure that analysing only a subsection of a sampler provides a reliable representation of the whole surface under real sampling conditions. Positional variability was evaluated using five predefined sections on a single sampler (3 × 1 cm each; Figure 2). To enable statistical assessment of within-sampler variation under realistic conditions, five sticky samplers collected from a domestic garden were used as biological replicates. DNA yields were quantified by qPCR, with each subsample analysed in technical triplicate.

**Figure 1.**
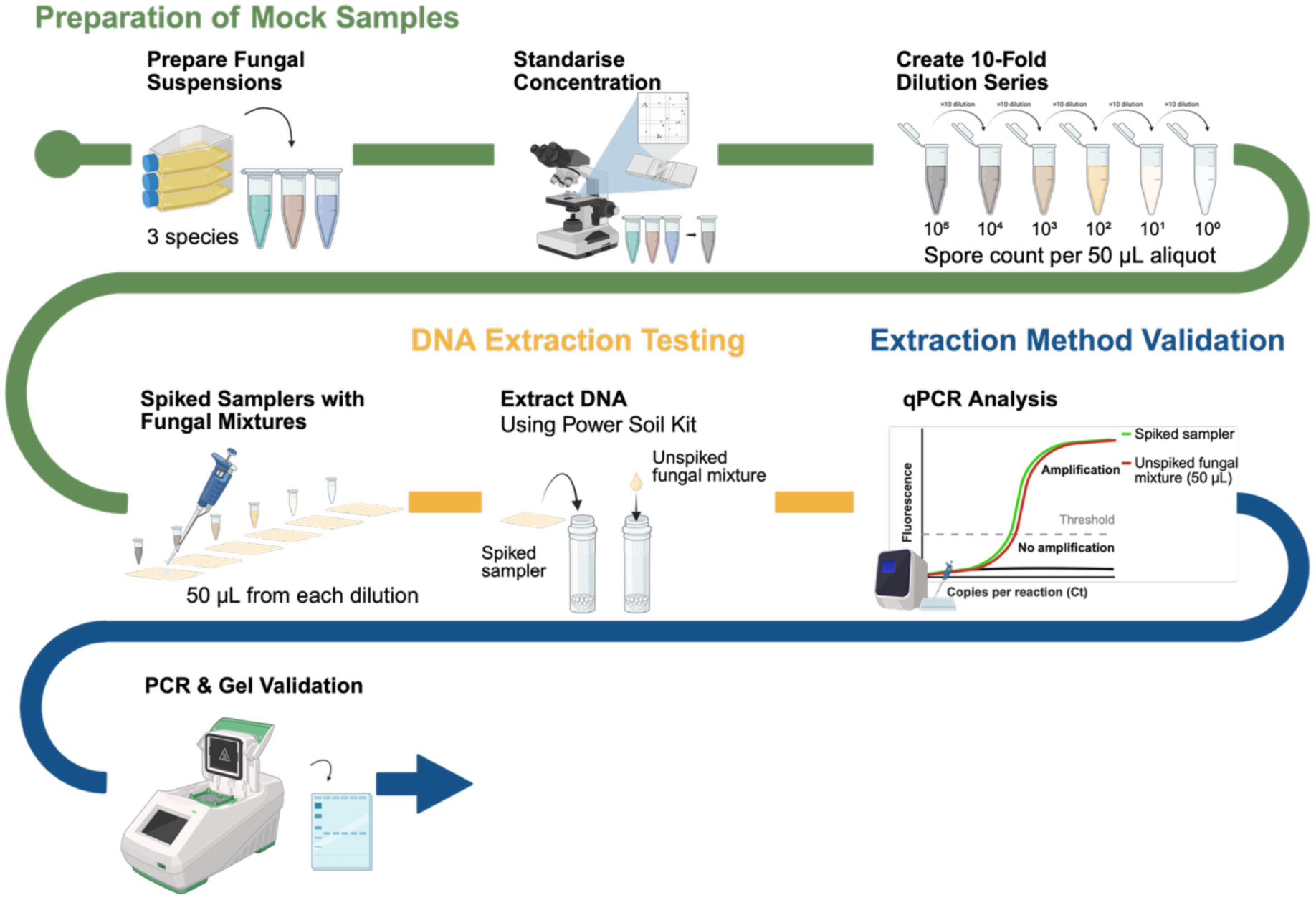
Workflow for the Mock Test, illustrating the preparation of a fungal mock community through a ten-fold serial dilution series (10⁵–10⁰ spores per 50 µL). This mock community was used in the optimisation and validation of the DNA-extraction protocol for the sticky samplers.

**Figure 2.**
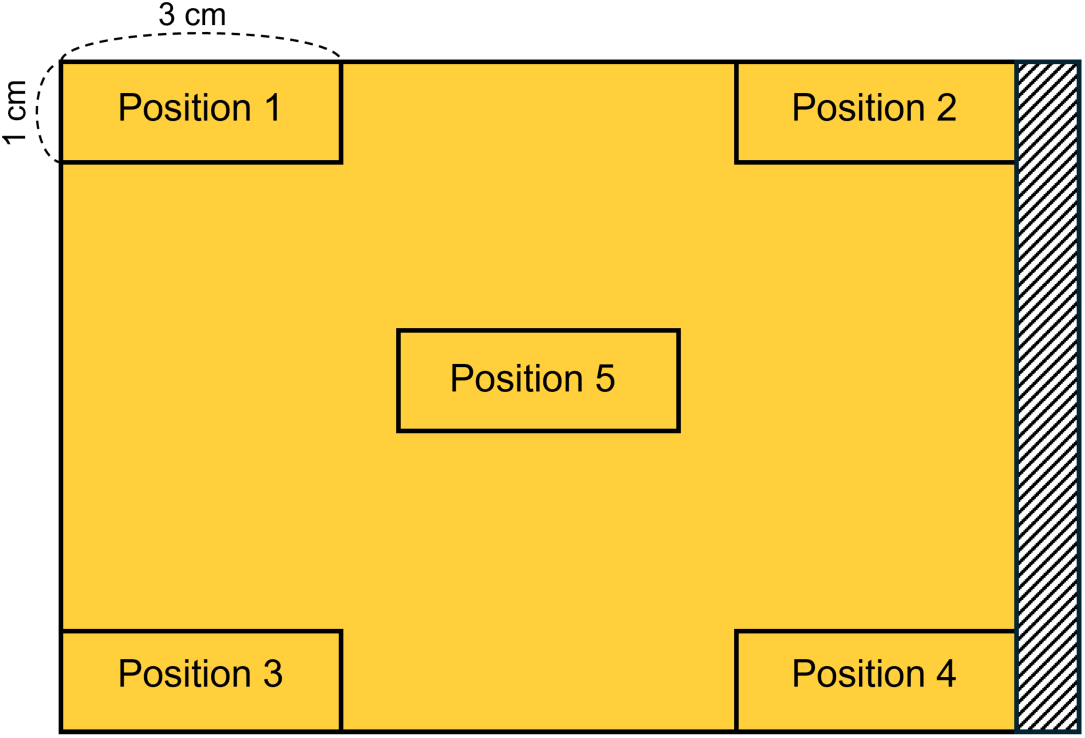
Sampling Layout of the Sticky Sampler, indicating positions 1–4 at the corners and position 5 at the centre. The hatched area on the right side represents a glue-free section.

### 2.4 Assessment of Sticky Sampler Performance and Suitability

#### 2.4.1 Baseline Assessment of Unused Sticky Sampler

Except for the mock test, in which the samplers were UV sterilised, all samplers were deployed without sterilisation to reflect realistic field conditions where pretreatment is often impractical. To evaluate whether non-UV-treated samplers carried background DNA, five unused sticky samplers (from separate, unopened packages), had neither been UV-treated nor exposed to the environment, were tested by qPCR. All reactions were conducted in technical triplicate to ensure reproducibility.

#### 2.4.2 Comparison with Other Glue-based Samplers

To assess the suitability of the sticky sampler material for downstream molecular analyses, we compared its DNA extraction efficiency with that of the black carbon tape described in the referenced study [19]. Comparisons were performed under two conditions: (i) by spiking both samplers with the mock fungal community used in the optimisation tests (Section 1), and (ii) by deploying them together in a domestic garden for four weeks, consistent with the exposure duration used in field sampling. DNA was extracted using the Direct extraction protocol described above and quantified by qPCR.

Additionally, a small-scale comparison was conducted to determine whether the performance of the sticky sampler (Microseal seals “B”, Bio-Rad) used in this study was specific to this product or comparable to other commercially available PCR sealing films. In addition to the Bio-Rad PCR sealing film, four other glue-based PCR sealing films were evaluated at the same sampling site under identical environmental conditions for four weeks (n = 2 per brand): Adhesive PCR film seals (VWR International, USA), SealPlate® film (Merck, USA), Easyseal™ Sealer, Clear (Greiner Bio-One, Austria), MicroAmp™ Optical Adhesive Film (Thermo Scientific, USA). Full product details are provided in Supplementary Table S1.

### 2.5 Time-Series Experiment

A 4-week deployment period was chosen because published studies employing the same sticky sampler have consistently used a 4-week sampling duration. To evaluate whether this deployment length is also suitable for DNA-based analysis, a time-series experiment was conducted over the same 4-week period. This experiment aimed to assess whether fungal DNA accumulation increases over time and whether a 4-week deployment provides sufficient DNA for downstream molecular analysis in both indoor and outdoor environments. At each indoor and outdoor location, four sticky samplers were deployed, with one retrieved at each of the following time points: 7, 14, 21 and 28 days. To obtain technical triplicates, three separate sections were subsampled from each sticky sampler, following the optimised DNA extraction method.

## Application of the Optimised Method to Environmental Samples

### 2.6 Application to Environmental Samples

The validated DNA extraction method was applied to samplers deployed under field conditions to evaluate its real-world applicability. Spatial variation in airborne fungal community was assessed using samplers collected from four agricultural farms in the UK, whereas seasonal variation was examined using samplers collected from a domestic garden across three seasons. In both cases, the sticky samplers were placed inside delta traps to protect them from environmental factors.

#### 2.6.1 Spatial Variation Study with Farm Samples

Samples were collected from four commercial farms located in different regions of England: Durham, Kent, Hampshire, and Hertfordshire (hereafter referred to as Farms A– D). At each farm, three spatially distinct sampling sites were selected (12 sites in total). Farm A cultivated two different crops, wheat and barley, in separate fields. Only the wheat field received fungicide treatment. In contrast, Farms B to D each grew a single crop: spring barley at Farm B and wheat at Farms C and D. Farm B was untreated, whereas Farms C and D were treated with fungicide. At each sampling site, a single sticky sampler was deployed. Three equal-sized subsamples were excised from each sampler and individually processed for DNA extraction, yielding a total of 36 DNA extracts (12 sites × 3 subsamples). Details of farm location, crop type, and fungicide treatment are summarised in Supplementary Table S2A.

#### 2.6.2 Seasonal Variation Study with Garden Samples

Samples were collected from a domestic garden in London during three seasons: autumn, winter, and spring (see Supplementary Table S2B). In each season, five sticky samplers were randomly selected as biological replicates, resulting in a total of 15 samples for downstream analysis.

#### 2.6.3 Analysing Fungal Community by ITS2 Metabarcoding

DNA from farms, garden, and control samples was PCR-amplified targeting the ITS2 region using ITS7F and ITS4R primers with Illumina adapters (see Table 2A), and amplification was verified by agarose gel electrophoresis (Supplementary Figure S6). Amplicons from farm (n = 36) and garden (n = 15) samples, as well as control samples (UV, NoUV, KC), were included in the sequencing run. All garden samples and control samples were sequenced in technical duplicate by loading the same PCR products into separate wells, and additional technical duplicates of selected farm samples were included to assess sequencing reproducibility, resulting in a total of 94 wells used for sequencing. Sequencing was performed on the Illumina MiSeq platform (version 3 chemistry, paired end 2 × 300 bp), generating approximately 15 million clusters per run at the Centre for Genomic Research, University of Liverpool.

**Table 2.**
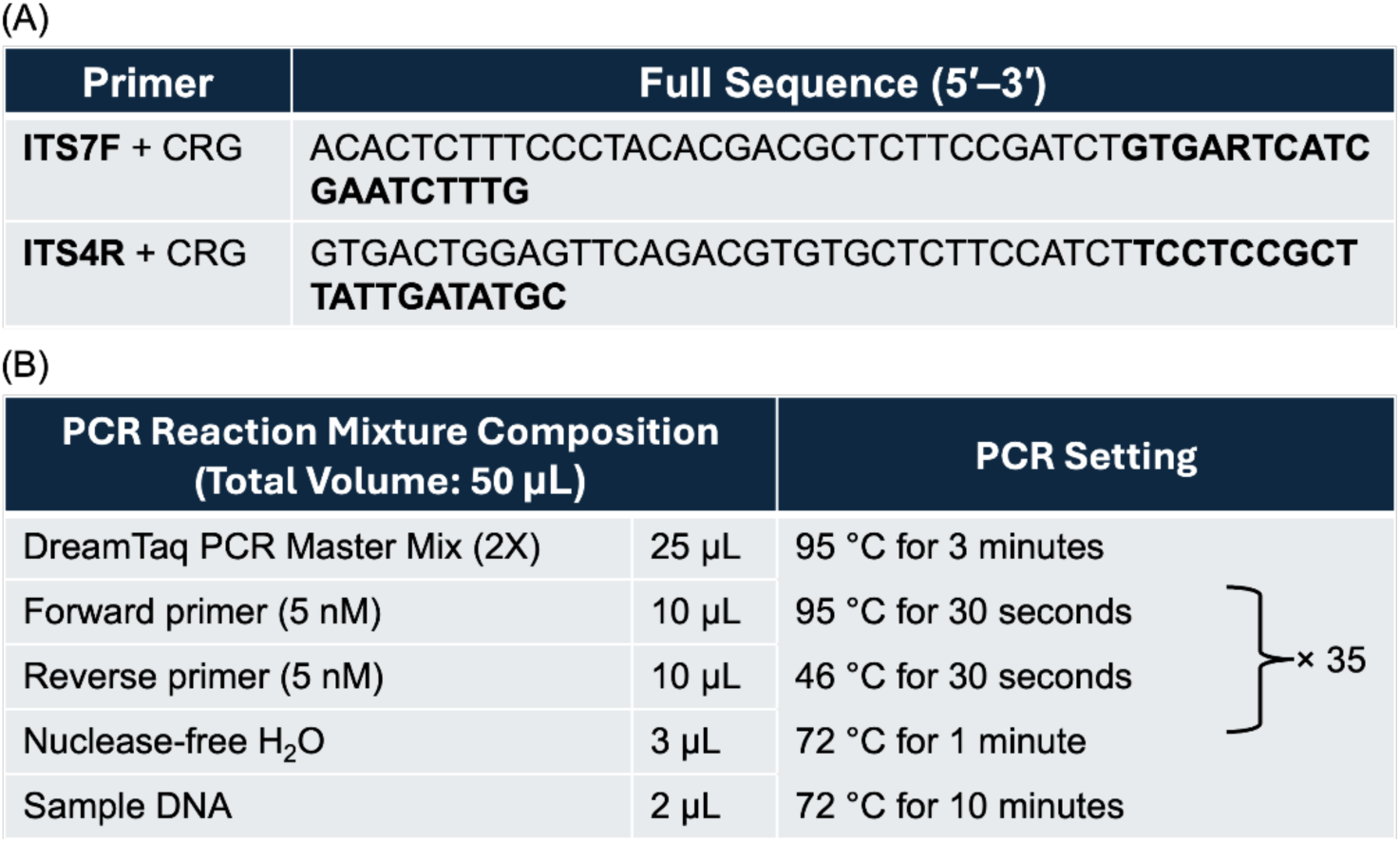
Primer Sequences and PCR Conditions Used for ITS2 Region Amplification in Sequencing Samples. (A) Primer sequences (5′–3′). The forward primer (ITS7F) was described by [23], and the reverse primer (ITS4R) by [24]. The full sequences include the core primer regions (in bold) with Illumina sequencing adaptors added by the Centre for Genomic Research, University of Liverpool. All primers were synthesised by Thermo Fisher Scientific, USA. (B) PCR reaction mixture composition and cycling conditions.

All analyses were conducted in R (version 4.4.3, released 28 February 2025) using RStudio. Amplicon sequence processing and denoising were performed using the DADA2 pipeline (version 1.36.0) [25], following the DADA2 ITS-specific workflow (version 1.8). Taxonomic classification was performed using the *assignTaxonomy* function against the UNITE reference database (General FASTA release version 10.0, released 19 February 2025) [26]. To ensure consistent and reliable classification, assignments were restricted to the genus level due to the limited resolution of ITS2 for species-level identification in many fungal taxa. Alpha diversity (within-sample diversity) was assessed using the Shannon index, and group-level differences were evaluated using Kruskal–Wallis tests with Dunn’s post hoc tests. Beta diversity (between-sample differences) was calculated based on Bray–Curtis dissimilarities. Ordination was performed via Principal Coordinates Analysis (PCoA) using the ordinate function in phyloseq (version 1.52.0), and variation in community composition was tested using PERMANOVA with 999 permutations.

## 3. Results

### 3.1 Direct Extraction Validation Comparison with Solvent-Based Approach

No meaningful amplification was detected with the Solvent-based method under any condition tested and this method was not pursued further. In contrast, the Direct DNA extraction method consistently yielded successful amplification across a wide concentration range (10⁵–10² spores per sample), demonstrating its feasibility for DNA downstream analysis.

## Sampling vs No Sampling Validation

To evaluate the detection limit of the direct extraction method, fungal mock samples containing 10⁵ to 10⁰ spores per sample were tested. Samples were either spores spiked onto sticky samplers before extraction (Sampling group) or spores extracted directly without sampler contact (No Sampling group; Figure 3A). Consistent amplification was observed in both groups down to 10² spores per sample, with no amplification detected at lower concentrations (Figure 3B). These results indicate that 10² spores per sample is the lowest concentration at which reliable amplification was consistently achieved under the tested conditions.

**Figure 3.**
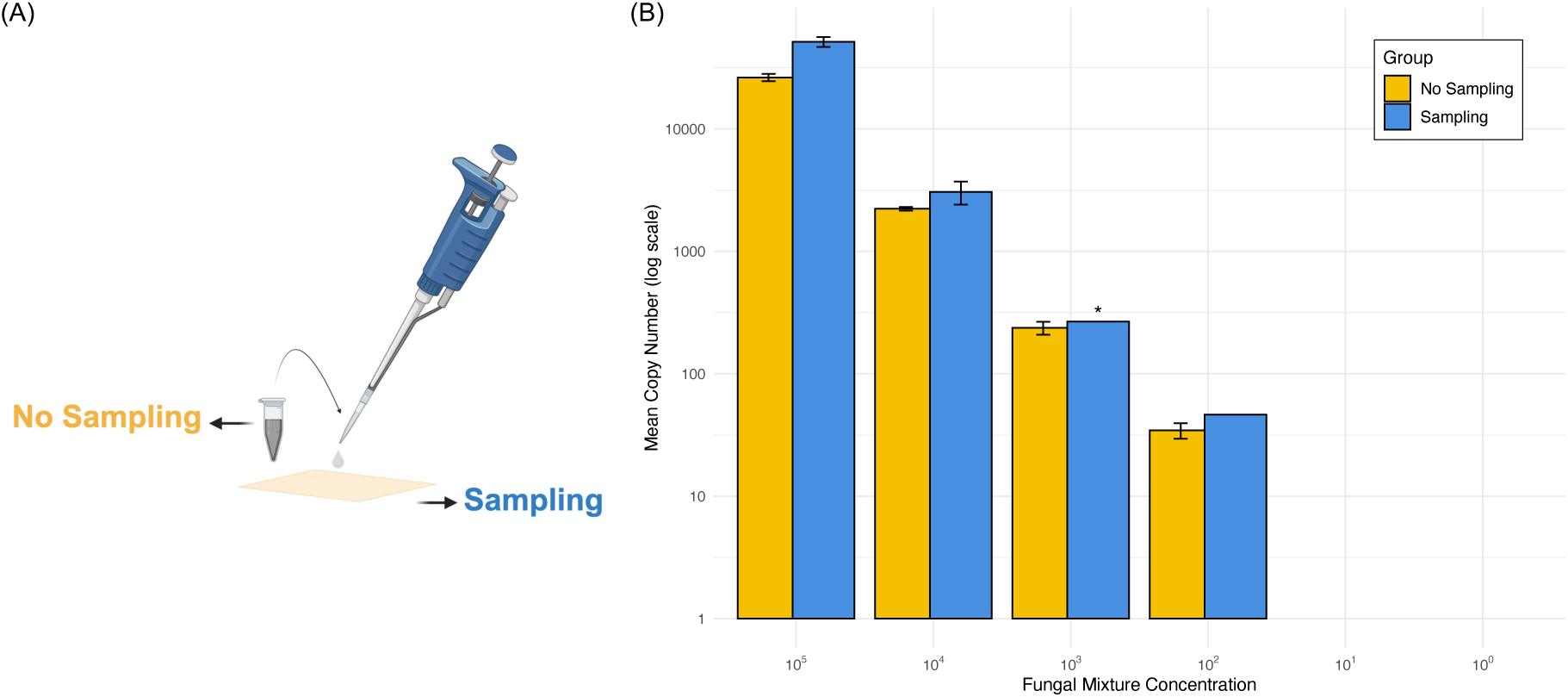
qPCR Results from Mock Test Using Direct DNA Extraction. (A) Illustration of the Sampling and No Sampling setup used in (B). (B) Comparison of DNA recovery between sticky samplers exposed to fungal mock mixtures (“Sampling”) and mock mixtures extracted directly without sampler contact (“No Sampling”). *Asterisk indicates detection in only one biological replicate. All samples in the mock test were prepared in duplicate (biological replicates) and analysed by qPCR in triplicate. qPCR èiciency = 69.94%, *R²* = 0.99.

Interestingly, the Sampling group showed slightly higher DNA copy numbers than the No Sampling group. At 10⁵ spores per sample, the Sampling group exhibited a mean DNA copy number of 51,477.06, nearly twice that of the No Sampling group (26,299.73). A similar pattern was observed at 10⁴ spores per sample, with mean values of 3,059.90 and 2,226.55, respectively. At the lowest concentration, 10² spores per sample, both groups yielded low but detectable quantities of DNA (46.42 and 34.48, respectively). At 10³ spores per sample, amplification was detected in only one of the two biological replicates in the Sampling group, resulting in the absence of a standard deviation bar. Nonetheless, the measured value (267.06) remained comparable to that of the No Sampling group (236.86). These quantitative measurements are provided in full in Supplementary Table S3. The corresponding NanoDrop-derived DNA concentration and purity measurements (A260/A280 and A260/A230 ratios) are provided in Supplementary Table S4.

## Positional Consistency of DNA Recovery

Fungal DNA copy numbers measured from five predefined positions (centre and four edges) across five independent samplers showed slight variation, but no consistent pattern suggesting positional bias was observed (Figure 4A). Although median values at certain edge positions (e.g., Edge2 and Edge4) were marginally higher, the overall variation remained within an acceptable range (Figure 4B). A one-way ANOVA suggested a weak trend towards positional differences (*F(*4, 20) = 2.626, *p* = 0.065), although the evidence was not strong enough to support statistical significance.

**Figure 4.**
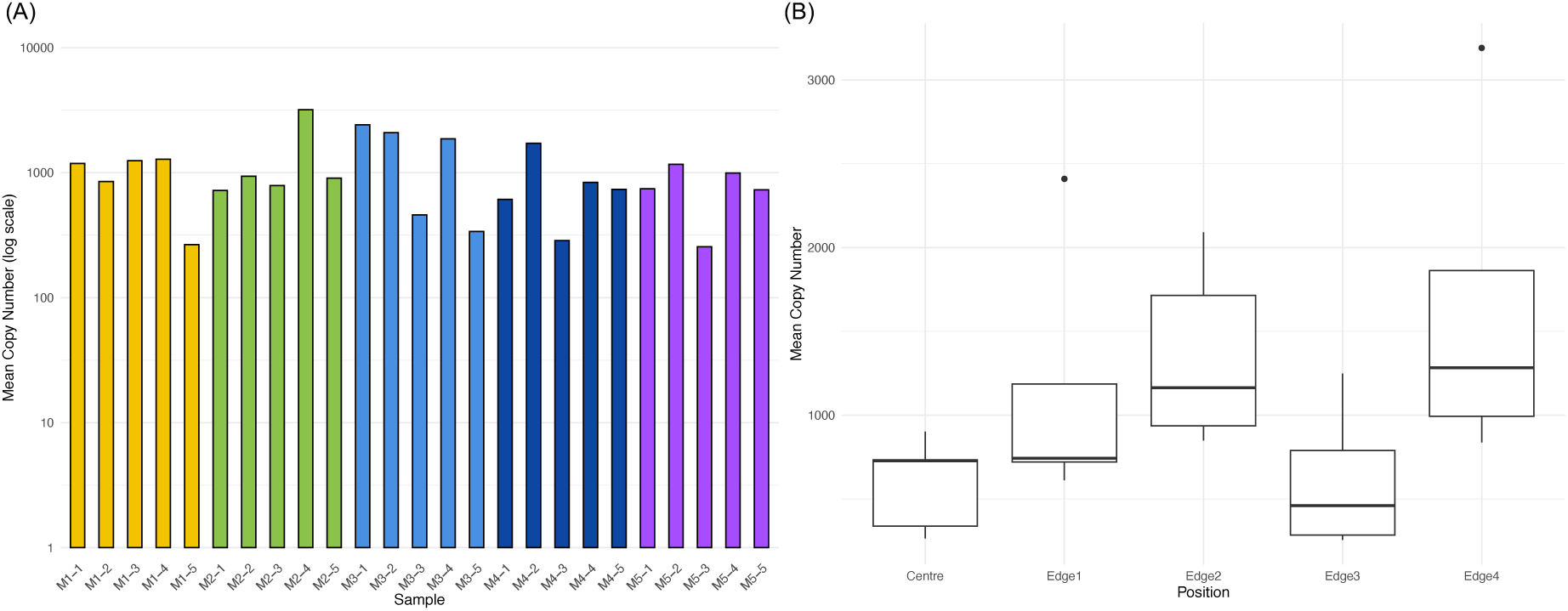
Positional Consistency of Fungal DNA Recovery from Sticky Samplers. (A) Mean fungal DNA copy number (log scale) measured at each sampling position on five sticky samplers (M = sampler number; positions 1–4 = corners, 5 = centre). Colours represent individual samplers. (B) Distribution of DNA copy number by sampling position. Boxplots show the median, interquartile range, and outliers.

Because baseline DNA recovery varied among samplers, a linear mixed-effects model (LMM) was fitted, treating sampling position as a fixed effect and sampler as a random effect. A small number of high-copy observations were present, but these occurred as isolated events and were not associated with specific sampling positions (Supplementary Figure S1). After accounting for sampler-to-sampler variation, the positional effect remained non-significant (LMM: *F* (4, 16) = 2.66, *p* = 0.071), indicating that the observed variation likely reflects occasional localised deposition rather than a systematic positional bias.

### 3.2 Sampler Performance and Suitability

#### Baseline DNA Yield of Sticky Sampler

No amplification was detected in two of the five unused, non-UV-treated sticky samplers. The remaining three showed very late amplification, with mean Ct values above 48, which was near the detection threshold (see Supplementary Table S5 for full Ct data). For reference, the 10⁰ standard, representing the lowest concentration point on the standard curve, yielded a Ct value of 38.36. These weak signals are likely attributable to background noise or non-specific amplification rather than true fungal DNA. The kit control exhibited similarly late amplification, whereas the NTC remained negative across all technical replicates.

#### Black Carbon Tape vs Sticky Sampler

In the comparison between the black carbon tape and the sticky sampler, no amplification was detected from either the mock fungal community-spiked or the field-exposed black tape samples. During extraction, the adhesive layer of the black tape detached and formed dark aggregates that appeared to interfere with DNA purification, whereas the sticky sampler remained physically intact and stable throughout the procedure.

#### Selected Sticky Sampler vs Other PCR Films

The additional small-scale comparison involving five PCR sealing films from four different manufacturers revealed that the Bio-Rad PCR sealing film (selected as sticky sampler in this study) yielded the highest DNA copy numbers under identical exposure and extraction conditions (Supplementary Figure S2). Although this supplementary test was limited in sample size (n = 2 per brand) and therefore did not allow for statistical analysis, the consistent trend supports the choice of the Bio-Rad film as the sticky sampler for further study.

#### Sequencing-Based Assessment of Background Contamination

Finally, sequencing-based assessment was also conducted to ensure the reliability of sequencing results and to evaluate potential contamination from the samplers. Amplicon sequence variants (ASVs), defined as unique sequences distinguished at single-nucleotide resolution following error modelling and denoising, were used as the unit of taxonomic resolution [25]. Non-UV-treated blanks, which were deployed for environmental sampling in the farms and the garden, contained six ASVs per sample with relatively higher read counts (85 and 132, respectively), whereas UV-treated blanks yielded only two ASVs with very low read counts (10 reads per sample; Supplementary Table S6A). Despite this, the overlap between ASVs in the non-UV-treated blanks and those in the experimental samples was minimal, 0.18% in garden samples and 0.13% in farm samples (detailed in Supplementary Table S6B). The kit control produced a single ASV with one read, and one technical replicate (KC-2) was excluded during the earlier quality filtering process. The taxonomic composition of control samples is shown in Supplementary Figure S3.

### 3.3 Time-Series Experiments: DNA Accumulation Over the Time

Fungal DNA accumulated over time in both indoor and outdoor conditions (Figure 5). Outdoor samplers consistently produced higher DNA copy numbers across the 4-week period, already yielding 512.4 copies on average at day 7 compared with 49.0 copies indoors. By day 7, DNA quantities in both environments had reached the qPCR detection limit defined by the mock validation experiment, although indoor values were close to the lower threshold. Across the full sampling duration, indoor yields ranged from 49.0 to 280.6 copies, whereas outdoor yields ranged from 512.4 to 2,991.9 copies, reflecting a markedly steeper rate of accumulation outdoors. Full quantitative values for all time points are provided in Supplementary Table S7.

**Figure 5.**
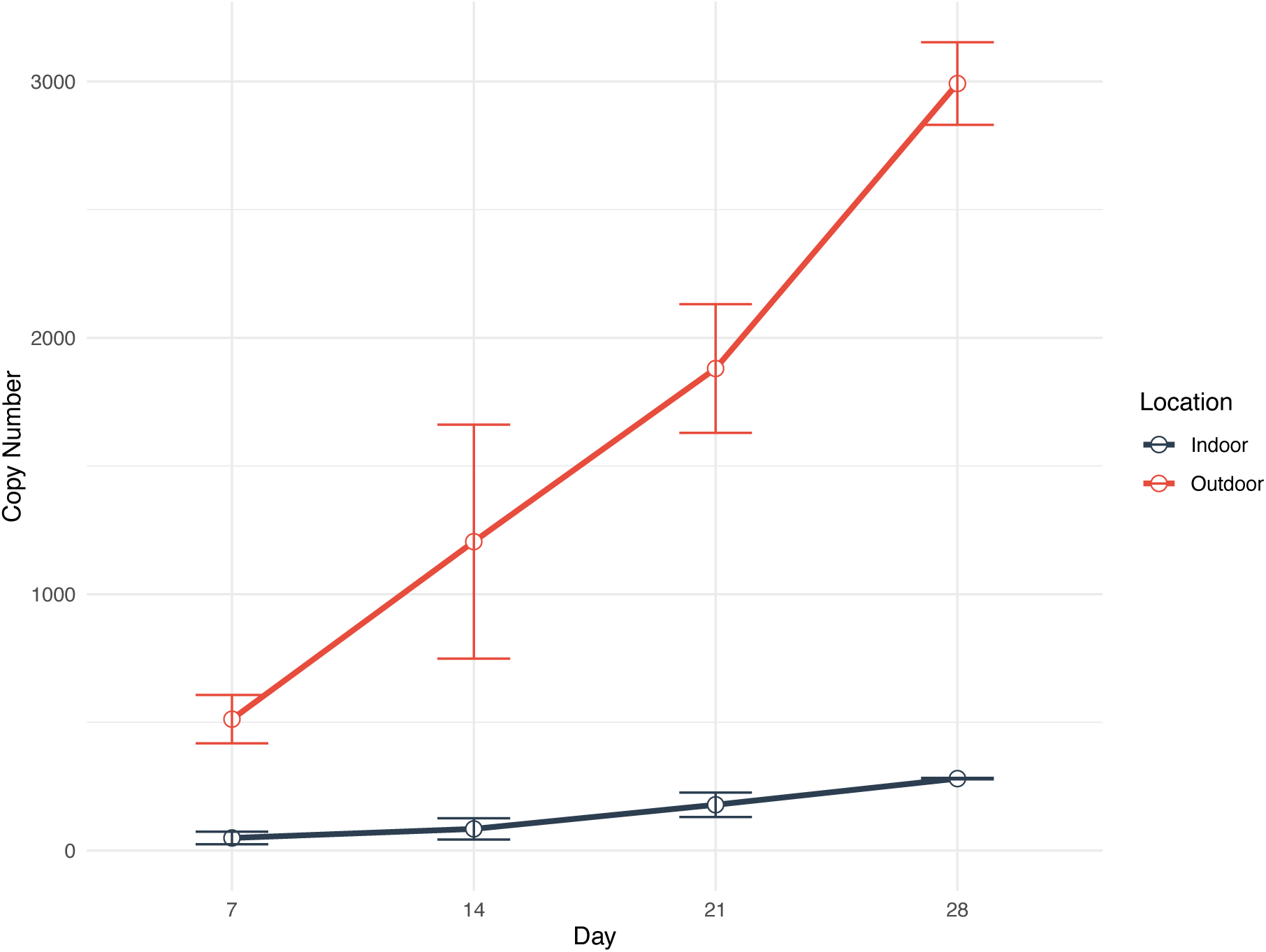
Temporal Variation in Fungal DNA Copy Numbers Detected from Sticky Samplers Exposed to Indoor and Outdoor Environments for 7, 14, 21, and 28 Days. Error bars represent the standard deviation (SD) among subsampling portions from the same sampler (n = 2–3 per location and time point).

### 3.4 Field Validation: Capturing Spatial and Seasonal Variation

Metadata for all environmental samples used in the ITS2 metabarcoding analysis, including both farm and garden datasets, are provided in Supplementary Table S8.

#### Sticky Samplers Capture Spatial Variation across Farms

Airborne fungal communities profiled by ITS2 metabarcoding exhibited clear spatial variation among the four farm sites. PCoA based on Bray–Curtis dissimilarities revealed distinct clustering by farm (Figure 6A) and this pattern was statistically supported by PERMANOVA (*R²* = 0.299, *F* = 7.69, *p* = 0.001). Notably, Farm A formed a completely distinct cluster with no overlap with the other farms (Figure 6A). Within-farm variation was also evident, as shown in Supplementary Figure S4, with PCoA revealing site-level clustering for individual farms and PERMANOVA confirming statistical significance (all *p* < 0.001).

**Figure 6.**
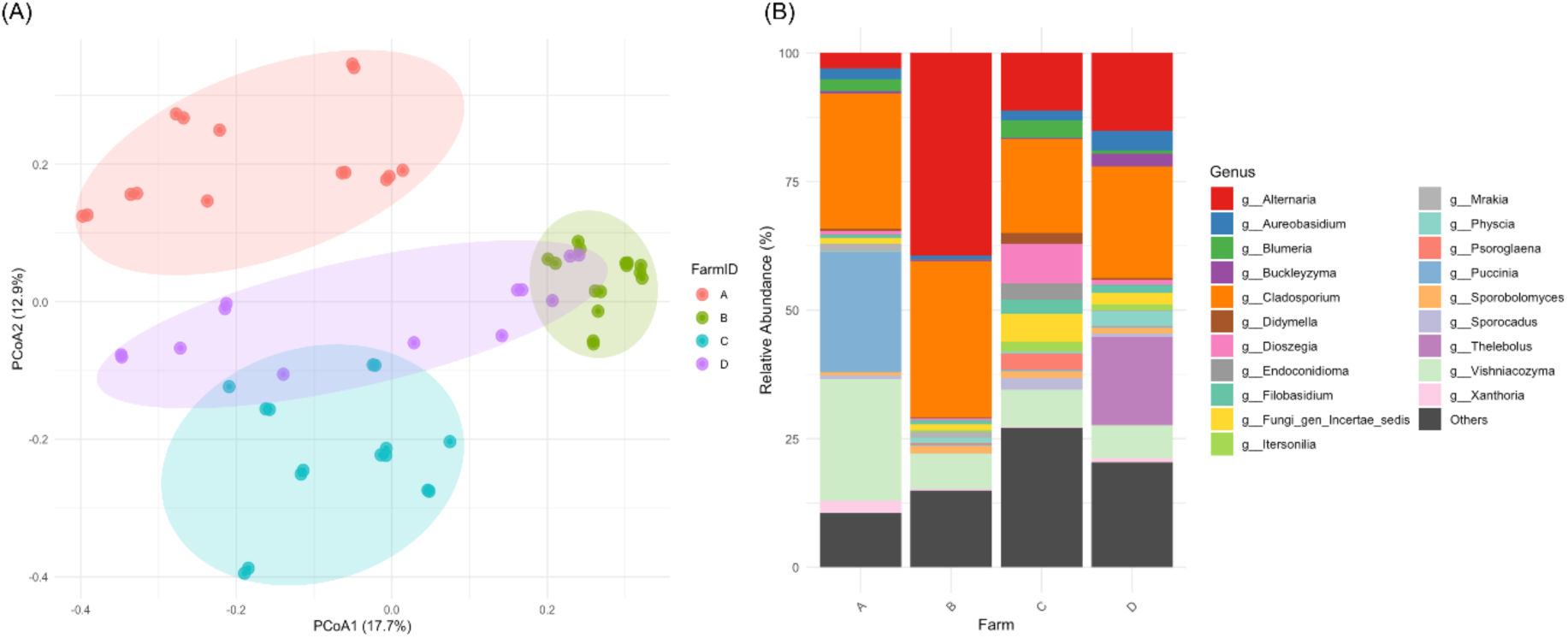
Community Variation in Airborne Fungal Composition across Farms. (A) PCoA showing distinct clustering by farm. Each point represents an individual sequencing library derived from farm samples (n = 58, Farms A–C = 15 each, Farm D = 13), coloured by farm identity. (B) Relative abundance of fungal genera across farms, based on the same set of samples, illustrating variation in taxonomic composition and the dominance of distinct genera at dìerent farms.

Genus-level fungal community composition differed substantially across farms (Figure 6B). Although *Cladosporium* was dominant across all farms, each farm exhibited distinct genus profiles. *Puccinia* and *Vishniacozyma* were particularly abundant in Farm A, while *Alternaria* was predominant in Farm B. Farms C and D showed a broader distribution of genera, with a higher proportion classified as ‘Others’ (genera outside the top 20) compared to Farms A and B. Within this broader distribution, particularly in Farm D, *Thelebolus* was distinctively present.

#### Sticky Samplers Capture Seasonal Dynamics in the Garden

Clear seasonal variation was evident in the garden fungal communities, with spring samples forming a distinct cluster from autumn and winter in ordination space (Figure 7B; PERMANOVA *R²* = 0.428, *F* = 10.11, *p* = 0.001). Diversity patterns mirrored these seasonal changes, with the Shannon index lowest in spring compared with winter (*p* = 0.049), while autumn and winter exhibited similar levels (*p* = 0.571; Figure 7A). However, the difference between spring and autumn did not reach statistical significance (*p* = 0.099; Figure 7A).

**Figure 7.**
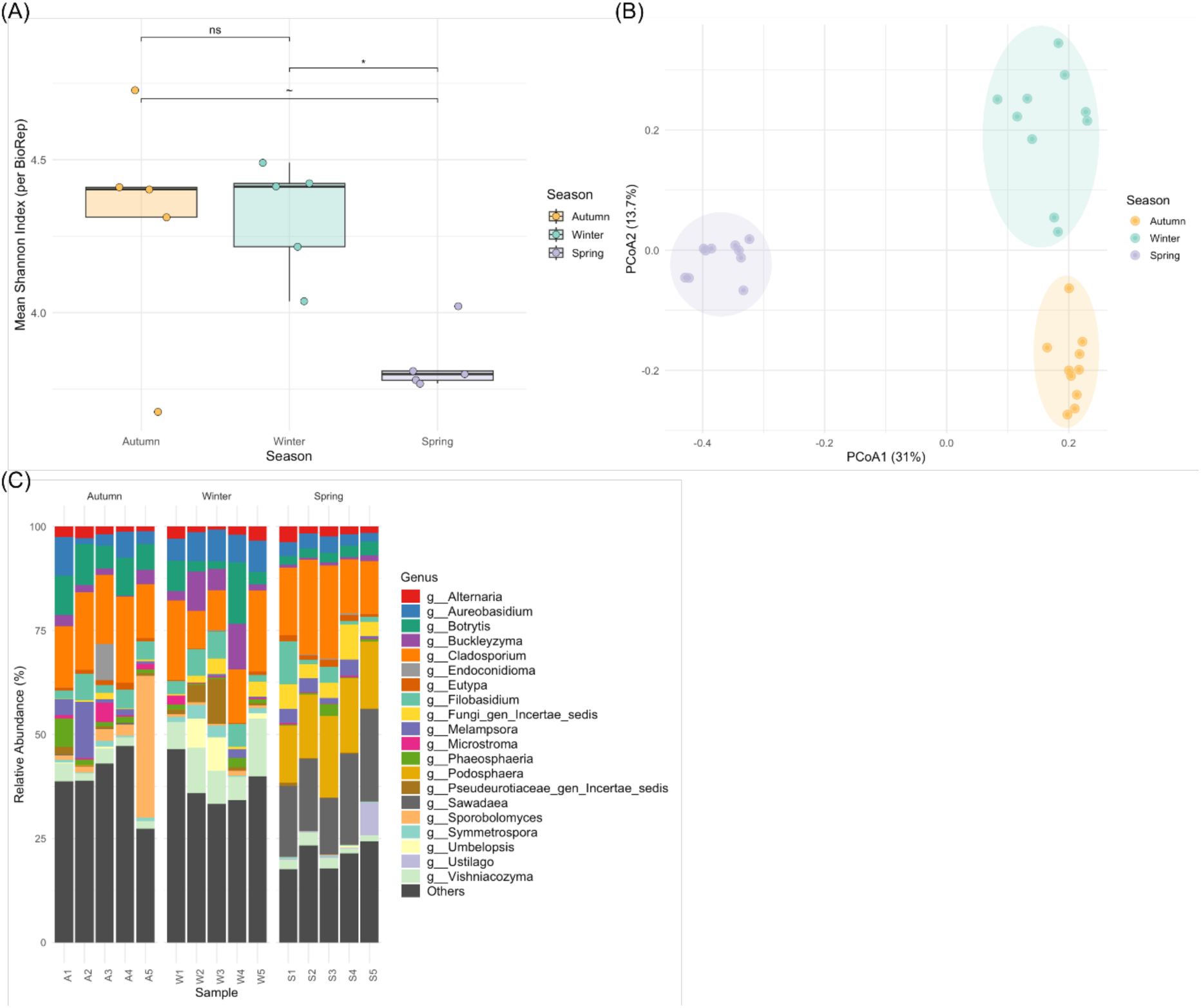
Seasonal Variation in Fungal Diversity in Garden Samples. (A) Shannon diversity across seasons, showing lower diversity in spring and higher in autumn and winter. Each point represents one biological replicate (n = 5 per season, mean of two technical replicates). Statistical significance is indicated as *p* < 0.05 (*), *p* < 0.1 (∼), and not significant (ns). (B) PCoA showing seasonal clustering. Each point represents an individual sequencing library (n = 10 per season). (C) Relative abundance of the top 20 fungal genera across three seasons, with remaining genera grouped as “Others”. Each bar represents one biological replicate.

Genus-level composition also varied seasonally, with distinct dominant taxa characterising each period (Figure 7C). *Podosphaera* and *Sawadaea* were more abundant in spring samples, whereas *Botrytis* occurred at low levels in spring but was more prominent in autumn and winter. *Aureobasidium* was consistently detected across seasons but appeared at higher relative abundance in winter. Autumn and winter samples showed greater genus-level diversity, indicated by higher proportions of genera grouped as “Others”. *Cladosporium* and *Alternaria* remained relatively stable across all seasons.

## 4. Discussion

This study introduces and validates an optimised DNA extraction workflow that enables direct ITS2 metabarcoding of fungal bioaerosols captured on glue-coated PCR sealing films. By eliminating solvent pretreatment and avoiding excessive degradation of the glue layer, the method achieves efficient and reproducible DNA recovery. When coupled with high-throughput sequencing, this approach enables the characterisation of both spatial and seasonal dynamics in airborne fungal communities. In addition, the workflow demonstrates the utility of glue-coated sealing films as a practical, cost-effective, and scalable platform for molecular surveillance of environmental mycobiomes.

The failure of Solvent-based extraction likely reflects detrimental interactions between solvents and the glue matrix, leading to DNA loss or inhibition during purification. In contrast, the Direct extraction method consistently yielded amplifiable DNA across a wide concentration range, demonstrating that glue dissolution is not required for efficient DNA recovery. Comparable DNA yields between the Sampling and No Sampling groups further support the robustness of the method. Interestingly, the Sampling group yielded slightly higher DNA copy numbers than the No Sampling group. This may be attributed to the central placement of the spiked mock community on the sticky sampler, which could have enhanced bead contact during lysis and improved cell disruption. These observations suggest that the sticky sampler, when paired with the optimised direct extraction method, may contribute to enhanced DNA recovery, rather than acting solely as a passive collector. Furthermore, the absence of positional bias indicates that analysing a portion of the sticky sampler is sufficient for a reliable estimate of the total fungal load. Together with the high DNA recovery rate, this positional consistency further supports the reliability of the optimised extraction method established in this study.

The material suitability of the sticky PCR sealing film was further verified for fungal bioaerosol monitoring. Baseline assessment of unused sticky samplers by qPCR revealed negligible background DNA, confirming their appropriateness for bioaerosol analysis without prior sterilisation. Consistently, sequencing-based validation of unused samplers further confirmed the reliability of the environmental datasets and showed negligible background contamination from the films (Supplementary Table S5). The Direct extraction approach developed in this study was conceptually inspired by a previously presented approach that employed black carbon tape as a glue-based sampler for bioaerosol sampling. However, when the same Direct extraction protocol was applied to black tape, DNA recovery consistently failed. During extraction, the formation of dark aggregates was observed, suggesting that the optimised lysis conditions may have been too harsh for the black tape, which may have led to adhesive degradation and loss of recoverable DNA. In contrast, the sticky PCR film retained residual adhesive after bead-beating, likely due to its glue composition designed for PCR sealing, which provides greater thermal and mechanical stability. These differences indicate that the extraction protocol established in this study is distinct from the reference work and specifically compatible with the sticky sampler material, supporting its selection as an effective platform for culture-independent fungal bioaerosol analysis.

In addition, comparison with other commercially available PCR sealing films further supported the effectiveness of the sticky sampler material. Among the films tested, the Bio-Rad film used in this study consistently produced higher DNA recovery than other brands (Supplementary Figure S2). This difference may reflect variations in the adhesive properties of the films among manufacturers, which in turn influence their compatibility with the extraction method. These findings demonstrate that not all PCR sealing films are equally compatible with the optimised direct extraction method, with the Bio-Rad film showing the highest performance under the conditions tested.

The time-series experiment showed that fungal DNA accumulated over time in both indoor and outdoor environments, demonstrating that sticky samplers function as passive collectors. However, interpretation should remain cautious because the experiment relied on a small sample size and a single indoor and outdoor sampling point. Outdoors, DNA yields exceeded the detection limit determined in the mock test at 7 days, yet no plateau was reached over the 4-week period, preventing estimation of a minimum sampling duration. Importantly, such a threshold must reflect not only qPCR detectability but also the DNA yield required for stable sequencing and community comparison. Indoors, DNA remained detectable at all time points but at consistently lower levels than outdoors. This trend agrees with previous studies [2, 27, 28], which report that airborne fungal levels are typically lower indoors than outdoors. However, the consistently low copy numbers raise the possibility that longer deployments are needed or that the culture-independent workflow is less effective under indoor conditions. Nonetheless, unpublished culture-based experiments using the same sampler have successfully recovered fungi from residential indoor environments. This suggests that the sampler is usable indoors and that the low yield observed here is more likely due to point-specific conditions or the limited sampling duration. To determine reliable minimum sampling times and generalisable indoor–outdoor contrasts, future time-series studies should expand the number and diversity of sampling locations and include buildings with varying ventilation characteristics.

The applicability of the sticky sampler combined with the optimised direct extraction method was further evaluated using farm and garden samples analysed by ITS2 metabarcoding. The workflow generated high-quality fungal community profiles from field-exposed samples, confirming its suitability for real-world bioaerosol monitoring beyond controlled laboratory conditions. The optimised workflow revealed clear spatial structuring of airborne fungal communities across farms (Figure 6B). Each farm harboured a significantly distinct community composition, reflecting site-specific environmental and management factors. In particular, the separation of Farm A in the PCoA appears to be associated with the markedly higher relative abundances of *Puccinia* and *Vishniacozyma*, which were far less dominant at other farms (Figure 6B). The geographical isolation of Farm A (Supplementary Figure S5), together with its unique crop composition and the fungicide treatment, may have influenced the prevalence of these taxa by providing distinct climatic conditions favourable to their growth [29, 30]. Farms B– D exhibited some degree of overlap, which may reflect their closer proximity and more comparable management practices relative to Farm A.

Moreover, underlying differences in soil fungal communities across the farms may also have contributed to the observed variation, as soil and plant debris often act as major sources of fungal spores released into the air through wind or agricultural disturbance [12, 31]. Beyond Farm A, genus-level profiles revealed additional distinctions among the remaining farms (Figure 6B). *Alternaria* was notably dominant in Farm B, possibly due to its fungicide-free status, whereas *Cladosporium* appeared consistently abundant across all farms regardless of crop type or management, in line with previous studies identifying it as a ubiquitous and environmentally resilient genus frequently detected in air samples [32]. Farms C and D, despite being geographically separate, shared similar profiles with a higher proportion of ‘Others’, which may indicate that fungicide use can alter competitive dynamics within the airborne mycobiota, potentially reducing dominant taxa and allowing secondary or resistant species to persist. Within these broader profiles, Farm D also showed a distinctive presence of *Thelebolus*, a genus associated with animal dung [33]. Although the exact source in this case is unknown due to limited contextual information, one possible explanation could be the influence of nearby livestock farming or the use of manure-based composts. Taken together, these results indicate that agricultural factors such as crop type and fungicide use may have a more pronounced impact on airborne fungal communities than geographic distance. Within-farm variation was also observed (Supplementary Figure S4), with differences in community composition among individual sites, supporting the idea that local microenvironments can influence fungal bioaerosols. However, site selection within farms was not standardised (e.g., entrance, centre, boundary), as sampling points were determined by farm owners, and precise coordinates cannot be revealed due to GDPR compliance. As a result, these within-farm differences should be interpreted as qualitative evidence of variation rather than as the basis for direct quantitative comparisons. To support more consistent within-and between-farm comparisons in future research, predefined spatial sampling criteria within farms should be considered, and within-farm biological replicates could be incorporated to better capture local heterogeneity.

Fungal community composition in garden samples showed clear seasonal variation, with the greatest dissimilarity observed between spring and winter (Figure 7A). This likely reflects a combination of factors, including substantial differences in temperature and humidity. By contrast, autumn and winter communities were more similar, which may be due to their temporal proximity and shared phenological stages of local vegetation, rather than strong climatic differentiation. Although the difference between spring and autumn was not statistically significant (*p* = 0.099), this may reflect greater variability within the autumn samples, which could have reduced the statistical power to detect seasonal divergence. PCoA ordination supported these patterns, with spring samples forming a distinct cluster (Figure 7B), likely driven by warmer and drier conditions known to influence spore dispersal and community composition. Shifts in genus-level composition mirrored the seasonal differences in diversity, with distinct dominant taxa characterising each period (Figure 7C). *Podosphaera* and *Sawadaea* were noticeably more abundant in spring samples. Both genera, belonging to the order Erysiphales, are foliar pathogens typically found on upper leaf surfaces during early plant growth stages [34, 35]. In contrast, *Botrytis*, a genus that thrives under cool and humid conditions [36], was present at low levels in spring but became more prominent in autumn and winter. Autumn and winter samples also showed greater genus-level diversity, likely reflecting the higher humidity and increased organic debris typical of these seasons. This was evident from the larger proportions of genera grouped as “Others”, representing rare or low-abundance taxa. *Aureobasidium* was detected in all seasons but appeared consistently and at higher abundance only in winter, suggesting a seasonal preference consistent with its known tolerance of cold and moist environments [35]. *Cladosporium* and *Alternaria* remained relatively stable in abundance across all seasons, underscoring their role as core components of the airborne mycobiome, in line with previous studies showing their persistence in urban air throughout the year [35, 37]. This pattern implies that passive sticky samplers are sensitive enough to detect seasonal transitions in airborne fungal emissions, from plant-associated foliar pathogens dominant in the warmer spring to saprotrophic taxa thriving under cooler and more humid conditions in autumn and winter, thereby capturing ecologically meaningful shifts in the airborne mycobiome.

Our findings demonstrate that sticky samplers, when paired with optimised DNA extraction and ITS2 metabarcoding, can effectively capture spatial and seasonal patterns in fungal bioaerosol communities. This practical workflow eliminates the need for adhesive removal while enabling fungal quantification and community-level characterisation from a single passive sampling device. The detection of groups such as *Fungi_gen_Incertae_sedis*, which represent sequences not confidently classified at the genus level, likely reflects current limitations in the ITS2 reference database rather than methodological error [38]. These unclassified groups may include poorly characterised or yet-undescribed fungal taxa. Taken together, these results highlight the value of sticky samplers as a practical and scalable tool for passive airborne mycobiome monitoring in both agricultural and urban settings. By providing a simple and effective molecular workflow, this study establishes a methodological foundation for future standardisation of glue-based bioaerosol sampling. The simplicity, low cost and ease of deployment of sticky samplers make them well suited for large-scale spatial and temporal monitoring programmes, including citizen-science initiatives. Nevertheless, future validation efforts should incorporate larger sample sizes and environmental samples collected across broader temporal and spatial gradients to further establish the reliability and generalisability of the adhesive film used in this study. Comparisons with established active air sampling techniques, such as cascade impactors and volumetric samplers, would also help define the applicability of the sticky sampler across different bioaerosol monitoring strategies.

## 5. Conclusion

This study demonstrates the feasibility and effectiveness of combining passive sticky samplers with an optimised DNA extraction and ITS2 metabarcoding workflow for culture-independent fungal bioaerosol profiling. The Direct DNA extraction workflow, applied to the adhesive surface of sticky samplers, enables efficient and reproducible recovery of fungal DNA, greatly simplifying bioaerosol processing and minimising sample loss. Validation with a mock community dilution series and field-collected samples confirmed its robustness across environments, revealing clear spatial and seasonal variation in airborne fungal assemblages. By linking methodological optimisation with ecological characterisation, this study establishes a practical and scalable framework for environmental mycobiome monitoring. Future work could integrate this approach into climate-coupled surveillance systems to advance large-scale monitoring of fungal bioaerosols in both indoor and outdoor environments and their environmental drivers.

## Availability of data and materials

All raw reads have been submitted to the National Centre for Biotechnology Information with accession number: PRJNA1355150.

## Code availability

All scripts can be found at https://github.com/AliceDay000/DADA2-ITS2-Metabarcoding-Pipeline/tree/main.

## Acknowledgements

This research was supported by a Natural Environment Research Council (NERC) NE/X00547X/1 and NE/X005259/1, Wellcome Trust 219551/Z/19/Z, and MRC MR/X020258/1. MCF is a fellow of the CIFAR Fungal Kingdoms programme.

## Author Contributions

M.C.F, R.L conceived the study, designed the sampling framework, and provided overall supervision and reviewed data analysis. J.J led the optimisation of the DNA extraction method, performed the experimental work, conducted data analysis, and drafted the manuscript. A.D developed the bioinformatic pipeline. R.L and M.C.F reviewed and approved the final manuscript. K.K., N.H., and H.D. coordinated farmer recruitment and carried out the deployment and retrieval of the sticky samplers at the farm sites.

