## Supplementary Material for "A Direct DNA Extraction Workflow for Metabarcoding Fungal Bioaerosols from Adhesive Samplers"

**Table S1 Commercially Available PCR Sealing Films Used for the Comparative Sampler Test**, including product name, manufacturer, country, and catalogue number.

| Product Name | Manufacturer | Country | Catalogue no. |
|---|---|---|---|
| MicroAmp™ Optical Adhesive Film | Thermo Scientific | USA | 4360954 |
| Adhesive PCR film seals | VWR | USA | 391-1254 |
| SealPlate® film | Merck | USA | Z369667-100EA |
| Easyseal™ Sealer, Clear | Greiner | Austria | 676001 |

**Table S2 Details of Environmental Samples** (A) shows details of the farm samples, including four agricultural sites (A-D) characterised by variation in location, crop type, and fungicide use. (B) shows details of the garden samples, including the seasonal sampling periods conducted over autumn, winter, and spring, each with a four-week sampling period.

(A)

| Farm | Location | Crop Type | Fungicide Applied |
|---|---|---|---|
| A | Durham | Winter Wheat | Azoxystrobin |
| | | Winter Barley | No fungicide applied |
| B | Kent | Spring Barley | No fungicide applied |
| C | Hampshire | Winter Wheat | Bixafen, Fluopyram |
| D | Hertfordshire | Winter Wheat | Tebuconazole |

(B)

| Season | Sampling Date (4 weeks) |
|---|---|
| Autumn | 2024 September – 2024 October |
| Winter | 2024 December – 2025 January |
| Spring | 2025 March – 2025 April |

**Table S3 qPCR Results from the Mock Test** The table reports Ct values and calculated DNA copy numbers for each dilution in the Sampling and No Sampling groups. Standard samples ($10^0$–$10^4$ DNA copies), blank control (not-exposed sampler), kit control (DNA extraction reagents only) and NTC were included. Efficiency (69.94%) and $R^2$ (0.99) for the standard curve are provided for reference.

| Group | Spore Concentration (spores/sample) | Ct Means | Copy Number | Note |
|---|---|---|---|---|
| Sampling | $10^5$ | 24.038 | 48118.76 | – |
| Sampling | $10^5$ | 23.792 | 54835.35 | – |
| Sampling | $10^4$ | 29.545 | 2596.38 | – |
| Sampling | $10^4$ | 28.969 | 3523.41 | – |
| Sampling | $10^3$ | 41.063 | 0.00 | – |
| Sampling | $10^3$ | 33.835 | 267.06 | – |
| Sampling | $10^2$ | 37.135 | 46.42 | – |

| | | | | |
|---|---|---|---|---|
| Sampling | $10^2$ | 37.135 | 46.42 | – |
| Sampling | $10^1$ | 40.970 | 0.00 | – |
| Sampling | $10^1$ | 41.821 | 0.00 | – |
| Sampling | $10^0$ | 40.159 | 0.00 | – |
| Sampling | $10^0$ | 42.685 | 0.00 | – |
| No Sampling | $10^5$ | 25.087 | 27597.01 | – |
| No Sampling | $10^5$ | 25.273 | 25002.45 | – |
| No Sampling | $10^4$ | 29.790 | 2280.03 | – |
| No Sampling | $10^4$ | 29.881 | 2173.07 | – |
| No Sampling | $10^3$ | 34.230 | 216.63 | – |
| No Sampling | $10^3$ | 33.907 | 257.09 | – |
| No Sampling | $10^2$ | 37.511 | 38.03 | – |
| No Sampling | $10^2$ | 37.901 | 30.93 | – |
| No Sampling | $10^1$ | 43.152 | 0.00 | – |
| No Sampling | $10^1$ | – | – | Omitted |
| No Sampling | $10^0$ | 46.239 | 0.00 | – |
| No Sampling | $10^0$ | Undetermined | Undetermined | – |
| $10^4$ | – | 20.378 | 18614.72 | Standard Sample |
| $10^3$ | – | 24.071 | 2627.13 | Standard Sample |
| $10^2$ | – | 28.967 | 195.93 | Standard Sample |
| $10^1$ | – | 33.724 | 15.74 | Standard Sample |
| $10^0$ | – | 37.218 | 2.47 | Standard Sample |
| BC | – | Undetermined | Undetermined | Blank Control |
| KC | – | Undetermined | Undetermined | Kit Control |
| NTC | – | Undetermined | Undetermined | No Template Control |

**Table S4 NanoDrop Results from the Mock Test** The table reports NanoDrop-derived DNA concentration (ng/μL), A260/A280, and A260/A230 ratios for DNA extracts from the Sampling and No Sampling groups used in the mock test.

| Group | Spore Concentration (spores/sample) | DNA Concentration (ng/μL) | A260/A280 | A260/A230 |
|---|---|---|---|---|
| Sampling | $10^5$ | 0.8 | 0.78 | 0.03 |
| Sampling | $10^5$ | 0.5 | 1.38 | 0.04 |
| Sampling | $10^4$ | 0.9 | 1.25 | 0.01 |
| Sampling | $10^4$ | 0.5 | 0.72 | 0.01 |
| Sampling | $10^3$ | 0.1 | 0.08 | 0.00 |
| Sampling | $10^3$ | 0.5 | 0.54 | 0.02 |

| | | | | |
|---|---|---|---|---|
| Sampling | $10^2$ | 2.1 | 1.00 | 0.08 |
| Sampling | $10^2$ | 4.9 | 1.30 | 0.30 |
| Sampling | $10^1$ | 5.3 | 1.12 | 0.14 |
| Sampling | $10^1$ | 0.5 | 0.86 | 0.01 |
| Sampling | $10^0$ | 7.0 | 0.44 | 0.05 |
| Sampling | $10^0$ | 0.9 | 0.72 | 0.02 |
| No Sampling | $10^5$ | 0.5 | 1.15 | 0.01 |
| No Sampling | $10^5$ | 19.9 | 1.39 | 0.37 |
| No Sampling | $10^4$ | 0.6 | 1.29 | 0.00 |
| No Sampling | $10^4$ | 3.5 | 1.57 | 0.01 |
| No Sampling | $10^3$ | 0.7 | 1.16 | 0.02 |
| No Sampling | $10^3$ | 8.7 | 1.46 | 0.13 |
| No Sampling | $10^2$ | 14.4 | 1.23 | 0.24 |
| No Sampling | $10^2$ | 2.1 | 0.96 | 0.03 |
| No Sampling | $10^1$ | 7.9 | 1.16 | 0.37 |
| No Sampling | $10^1$ | 4.0 | 1.15 | 0.02 |
| No Sampling | $10^0$ | 0.6 | 0.69 | 0.01 |
| No Sampling | $10^0$ | 2.7 | 1.36 | 0.00 |

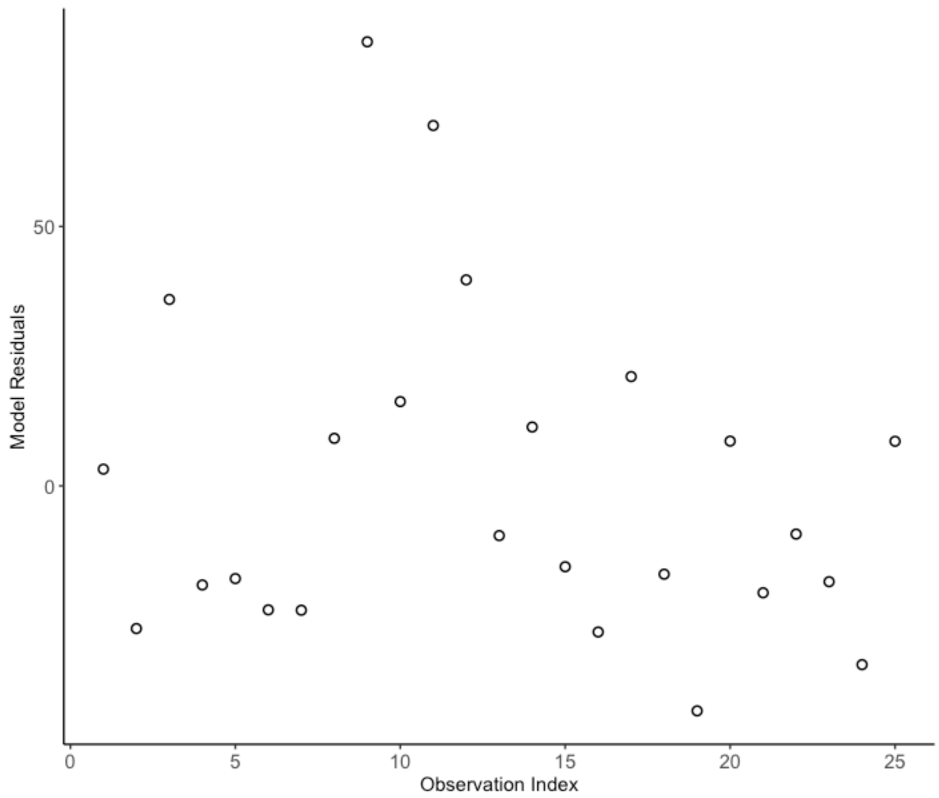


**Figure S1 Residual Plot from the Linear Mixed-Effects Model Assessing Positional Effects**
Model residuals (observed–predicted differences) were evenly distributed without positional

structure. A small number of high-copy observations occurred independently of sampling position and did not affect the overall conclusion.

**Table S5 Baseline qPCR Results from Unused, Non-UV-Treated Sticky Samplers** Ct values for five unused samplers (B1-B5), along with the kit control (BKC) and no-template control (NTC), are presented. Two samples (B1 and B3) showed no amplification, while the remaining three displayed very late amplification (Ct > 48). The $10^0$ standard yielded a Ct value of 38.36 (data not shown), indicating that amplification in these samples occurred beyond the quantifiable range.

| Sample | Replicate 1 | Replicate 2 | Replicate 3 | CT Mean | Note |
|---|---|---|---|---|---|
| B1 | – | – | – | – | No Amplification |
| B2 | 48.805 | – | 48.424 | 48.615 | Late CT |
| B3 | – | – | – | – | No Amplification |
| B4 | 49.731 | 49.225 | – | 49.478 | Late CT |
| B5 | – | – | 48.017 | 48.017 | Late CT |
| BKC | 47.471 | 48.865 | 47.885 | 48.073 | Kit Control |
| NTC | – | – | – | – | No Template Control |

**Table S6 Sequencing Summary and Contamination Assessment for Control Samples (**A) Total reads and observed ASVs in UV-treated blanks (UV), non-UV-treated blanks (NoUV), and kit controls (KC). The numbers following the dash indicate technical replicates derived from the same source sample. (B) Mean ASV richness (±SD) in garden and farm samples, and the number and percentage of ASVs shared with NoUV.

(A)

| Sample | Sample Type | Total Reads | Observed ASVs |
|---|---|---|---|
| UV-1 | UV-treated blank control | 10 | 2 |
| UV-2 | | 10 | 2 |
| NoUV-1 | Non-UV-treated blank control | 85 | 6 |
| NoUV-2 | | 132 | 6 |
| KC-1 | Kit control | 1 | 1 |
| KC-2 | | – | – |

(B)

| Sample Group | Mean ASV ± SD | Shared ASVs with NoUV | Shared (%) |
|---|---|---|---|
| NoUV (n=2) | 6.0 ± 0.0 | – | – |
| Garden (n=30) | 369.2 ± 117.0 | 6 | 0.18% |
| Farm (n=58) | 216.3 ± 62.9 | 4 | 0.13% |

**Table S7 qPCR Quantification of Fungal DNA in the Indoor and Outdoor Time-Series Sampling Experiment** The table reports DNA copy numbers for each time point, including standard samples ($10^0$-$10^4$ DNA copies), blank control (not-exposed sampler), kit control (DNA extraction reagents only) and NTCs. Replicate numbers correspond to three subsampling portions taken from each sampler. Each extract was analysed in qPCR technical triplicate. Efficiency (69.00%) and $R^2$ (0.98) for the standard curve are provided for reference.

| Sample | Location | Day | Replicate | Ct Mean | Copy Number | Note |
|---|---|---|---|---|---|---|
| ID1 | Indoor | 7 | 1 | 40.721 | 77.37 | – |
| ID1 | Indoor | 7 | 2 | 42.294 | 33.90 | – |
| ID1 | Indoor | 7 | 3 | 42.201 | 35.59 | – |
| ID2 | Indoor | 14 | 1 | 41.187 | 60.58 | – |
| ID2 | Indoor | 14 | 2 | 41.202 | 60.11 | – |
| ID2 | Indoor | 14 | 3 | 39.700 | 132.22 | – |
| ID3 | Indoor | 21 | 1 | 39.526 | 144.88 | – |
| ID3 | Indoor | 21 | 2 | 38.800 | 212.03 | – |
| ID3 | Indoor | 21 | 3 | – | – | Omitted |
| ID4 | Indoor | 28 | 1 | 38.280 | 278.51 | – |
| ID4 | Indoor | 28 | 2 | – | – | Omitted |
| ID4 | Indoor | 28 | 3 | 38.252 | 282.62 | – |
| OD1 | Outdoor | 7 | 1 | 36.885 | 579.02 | – |
| OD1 | Outdoor | 7 | 2 | – | – | Omitted |
| OD1 | Outdoor | 7 | 3 | 37.384 | 445.68 | – |
| OD2 | Outdoor | 14 | 1 | 34.798 | 1731.16 | – |
| OD2 | Outdoor | 14 | 2 | 35.990 | 926.24 | – |
| OD2 | Outdoor | 14 | 3 | 35.927 | 957.19 | – |
| OD3 | Outdoor | 21 | 1 | 34.379 | 2157.34 | – |
| OD3 | Outdoor | 21 | 2 | 34.869 | 1667.68 | – |
| OD3 | Outdoor | 21 | 3 | 34.707 | 1815.87 | – |
| OD4 | Outdoor | 28 | 1 | 33.867 | 2822.25 | – |
| OD4 | Outdoor | 28 | 2 | 33.743 | 3010.74 | – |
| OD4 | Outdoor | 28 | 3 | 33.662 | 3142.79 | – |
| $10^4$ | – | – | – | 24.344 | 23195.97 | Standard Sample |
| $10^3$ | – | – | – | 29.271 | 1748.28 | Standard Sample |
| $10^2$ | – | – | – | 33.428 | 197.41 | Standard Sample |
| $10^1$ | – | – | – | 38.100 | 17.01 | Standard Sample |
| $10^0$ | – | – | – | 41.871 | 2.35 | Standard Sample |
| BC | – | – | – | 47.321 | 0.00 | Blank Control |
| KC | – | – | – | Undetermined | Undetermined | Kit Control |

| | | | | | | |
|---|---|---|---|---|---|---|
| NTC | – | – | – | Undetermined | Undetermined | No Template Control |

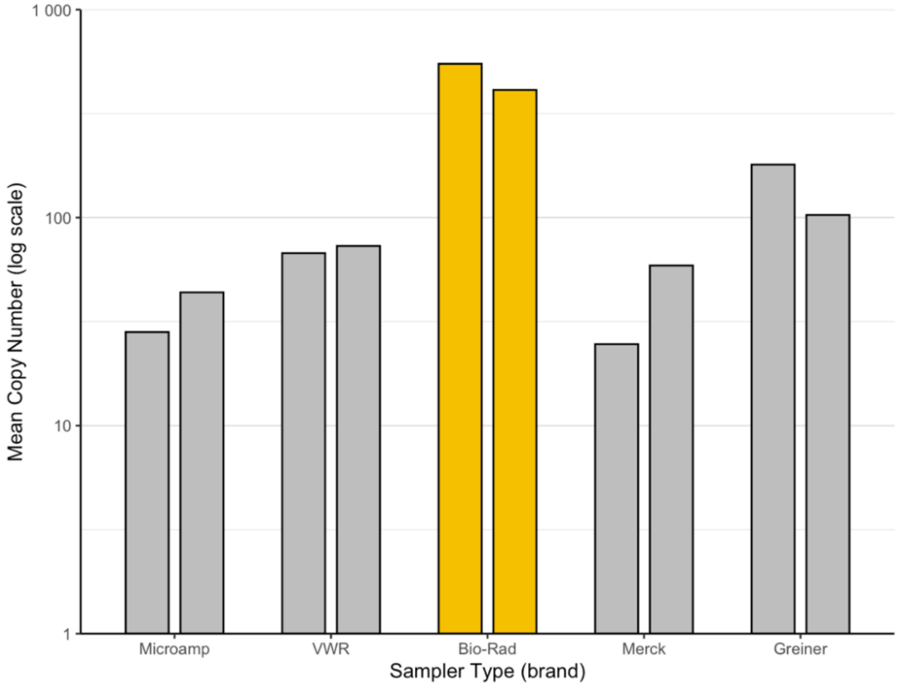


**Figure S2 Comparison of Mean Fungal DNA Copy Numbers Recovered from Five Different Glue-Based PCR Sealing Films** Films from five commercial brands (Microamp, VWR, Bio-Rad, Merck, and Greiner) were deployed for four weeks under identical environmental conditions. Each bar represents the mean DNA copy number obtained from two samplers per brand. This small-scale comparison supported the suitability of the Bio-Rad film, which served as the sticky sampler in this study.

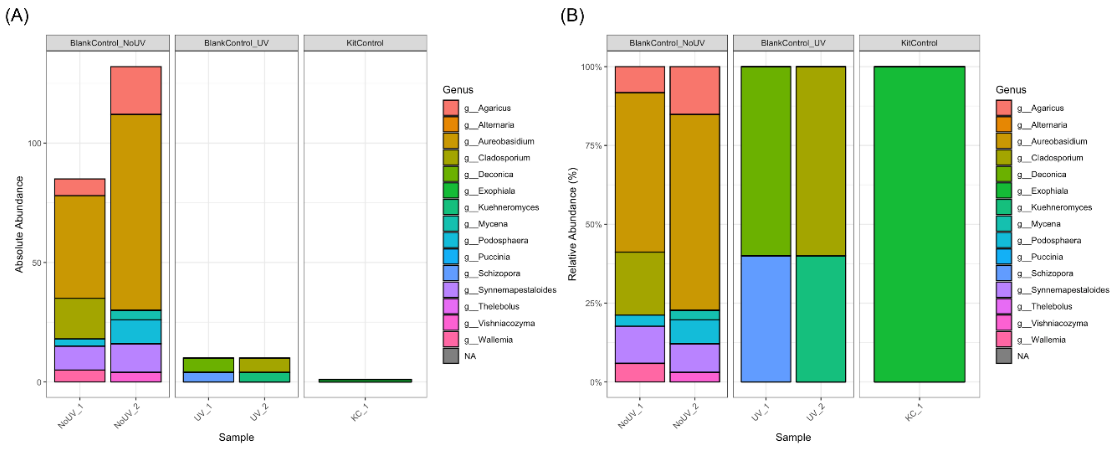


**Figure S3 Genus-Level Fungal Composition of Blank and Kit Control Samples** (A) Absolute abundance of fungal genera of control samples (B) Relative abundance of fungal genera

**Table S8 Metadata for Farm, Garden, and Control Samples Used in ITS2 Metabarcoding Analysis** The dataset includes farm, garden, and control samples (KCs, UVs, and No-UVs). All Sample IDs begin with a numeric prefix, followed by an alphabetic code that specifies the sample category. IDs containing “F” directly after the prefix correspond to farm samples: the letter following “F” denotes the farm (A–D), the number specifies the within-farm site (1–3), and the digit after the final dash indicates the subsample (e.g., 34-FA1-2 = Farm A, Site 1, Subsample 2). Farm Site values (1–3) represent three independent sampling points within each farm; their exact locations differ by farm because the three sampling sites were selected by farm owners. IDs in which the first alphabetic character is not “F” correspond to garden samples, where the initial letter indicates the sampling season (“S” = Spring, “A” = Autumn, “W” = Winter).

| Sample ID | Sample Type | Farm ID | Farm Site | TechRep | TechRep Group | BioRep | BioRep Group | SeqRep |
|---|---|---|---|---|---|---|---|---|
| 01-A1 | Garden | NA | NA | NA | A1 | 1 | A | 1 |
| 02-S3 | Garden | NA | NA | NA | S3 | 3 | S | 1 |
| 03-FB1-1 | Farm | B | 1 | 1 | FB1 | NA | NA | 1 |
| 04-FC2-1 | Farm | C | 2 | 1 | FC2 | NA | NA | 1 |
| 05-FD3-1 | Farm | D | 3 | 1 | FD3 | NA | NA | 1 |
| 06-W2 | Garden | NA | NA | NA | W2 | 2 | W | 1 |
| 07-FA2-2 | Farm | A | 2 | 2 | FA2 | NA | NA | 1 |
| 08-FC2-2 | Farm | C | 2 | 2 | FC2 | NA | NA | 1 |
| 09-A2 | Garden | NA | NA | NA | A2 | 2 | A | 1 |
| 10-S4 | Garden | NA | NA | NA | S4 | 4 | S | 1 |
| 11-FB1-2 | Farm | B | 1 | 2 | FB1 | NA | NA | 1 |
| 12-FC2-2 | Farm | C | 2 | 2 | FC2 | NA | NA | 2 |
| 13-FD3-2 | Farm | D | 3 | 2 | FD3 | NA | NA | 1 |
| 14-W3 | Garden | NA | NA | NA | W3 | 3 | W | 1 |
| 15-FA3-1 | Farm | A | 3 | 1 | FA3 | NA | NA | 1 |
| 16-FC3-1 | Farm | C | 3 | 1 | FC3 | NA | NA | 1 |
| 17-A3 | Garden | NA | NA | NA | A3 | 3 | A | 1 |
| 18-S5 | Garden | NA | NA | NA | S5 | 5 | S | 1 |
| 19-FB1-3 | Farm | B | 1 | 3 | FB1 | NA | NA | 1 |
| 20-FC2-3 | Farm | C | 2 | 3 | FC2 | NA | NA | 1 |
| 21-FD3-3 | Farm | D | 3 | 3 | FD3 | NA | NA | 1 |
| 22-W4 | Garden | NA | NA | NA | W4 | 4 | W | 1 |
| 23-FA3-2 | Farm | A | 3 | 2 | FA3 | NA | NA | 1 |
| 24-FC3-2 | Farm | C | 3 | 2 | FC3 | NA | NA | 1 |
| 25-A4 | Garden | NA | NA | NA | A4 | 4 | A | 1 |
| 26-FA1-1 | Farm | A | 1 | 1 | FA1 | NA | NA | 1 |
| 27-FB2-1 | Farm | B | 2 | 1 | FB2 | NA | NA | 1 |
| 28-FC3-1 | Farm | C | 3 | 1 | FC3 | NA | NA | 2 |

| | | | | | | | | |
|---|---|---|---|---|---|---|---|---|
| 29-UV | Blank Control _UV | NA | NA | NA | UV | NA | NA | 1 |
| 30-W5 | Garden | NA | NA | NA | W5 | 5 | W | 1 |
| 31-FB1-1 | Farm | B | 1 | 1 | FB1 | NA | NA | 2 |
| 32-FD1-1 | Farm | D | 1 | 1 | FD1 | NA | NA | 1 |
| 33-A5 | Garden | NA | NA | NA | A5 | 5 | A | 1 |
| 34-FA1-2 | Farm | A | 1 | 2 | FA1 | NA | NA | 1 |
| 35-FB2-2 | Farm | B | 2 | 2 | FB2 | NA | NA | 1 |
| 36-FC3-2 | Farm | C | 3 | 2 | FC3 | NA | NA | 2 |
| 37-NoUV | Blank Control _NoUV | NA | NA | NA | NoUV | NA | NA | 1 |
| 38-S1 | Garden | NA | NA | NA | S1 | 1 | S | 1 |
| 39-FB1-2 | Farm | B | 1 | 2 | FB1 | NA | NA | 2 |
| 40-FD1-2 | Farm | D | 1 | 2 | FD1 | NA | NA | 1 |
| 41-W1 | Garden | NA | NA | NA | W1 | 1 | W | 1 |
| 42-FA1-3 | Farm | A | 1 | 3 | FA1 | NA | NA | 1 |
| 43-FB2-3 | Farm | B | 2 | 3 | FB2 | NA | NA | 1 |
| 44-FC3-3 | Farm | C | 3 | 3 | FC3 | NA | NA | 1 |
| 45-KC | Kit Control | NA | NA | NA | KC | NA | NA | 1 |
| 46-S2 | Garden | NA | NA | NA | S2 | 2 | S | 1 |
| 47-FB2-1 | Farm | B | 2 | 1 | FB2 | NA | NA | 2 |
| 48-FD2-1 | Farm | D | 2 | 1 | FD2 | NA | NA | 1 |
| 49-W2 | Garden | NA | NA | NA | W2 | 2 | W | 2 |
| 50-FA2-1 | Farm | A | 2 | 1 | FA2 | NA | NA | 1 |
| 51-FB3-1 | Farm | B | 3 | 1 | FB3 | NA | NA | 1 |
| 52-FD1-1 | Farm | D | 1 | 1 | FD1 | NA | NA | 2 |
| 53-A1 | Garden | NA | NA | NA | A1 | 1 | A | 2 |
| 54-S3 | Garden | NA | NA | NA | S3 | 3 | S | 2 |
| 55-FB2-2 | Farm | B | 2 | 2 | FB2 | NA | NA | 2 |
| 56-FD2-2 | Farm | D | 2 | 2 | FD2 | NA | NA | 1 |
| 57-W3 | Garden | NA | NA | NA | W3 | 3 | W | 2 |
| 58-FA2-2 | Farm | A | 2 | 2 | FA2 | NA | NA | 2 |
| 59-FB3-2 | Farm | B | 3 | 2 | FB3 | NA | NA | 1 |
| 60-FD1-2 | Farm | D | 1 | 2 | FD1 | NA | NA | 2 |
| 61-A2 | Garden | NA | NA | NA | A2 | 2 | A | 2 |
| 62-S4 | Garden | NA | NA | NA | S4 | 4 | S | 2 |

| | | | | | | | | |
|---|---|---|---|---|---|---|---|---|
| 63-FB3-1 | Farm | B | 3 | 1 | FB3 | NA | NA | 2 |
| 64-UV | Blank Control _UV | NA | NA | NA | UV | NA | NA | 2 |
| 65-W4 | Garden | NA | NA | NA | W4 | 4 | W | 2 |
| 66-FA2-3 | Farm | A | 2 | 3 | FA2 | NA | NA | 1 |
| 67-FB3-3 | Farm | B | 3 | 3 | FB3 | NA | NA | 1 |
| 68-FD1-3 | Farm | D | 1 | 3 | FD1 | NA | NA | 1 |
| 69-A3 | Garden | NA | NA | NA | A3 | 3 | A | 2 |
| 70-S5 | Garden | NA | NA | NA | S5 | 5 | S | 2 |
| 71-FB3-2 | Farm | B | 3 | 2 | FB3 | NA | NA | 2 |
| 72-NoUV | Blank Control _NoUV | NA | NA | NA | NoUV | NA | NA | 2 |
| 73-W5 | Garden | NA | NA | NA | W5 | 5 | W | 2 |
| 74-FA3-1 | Farm | A | 3 | 1 | FA3 | NA | NA | 2 |
| 75-FC1-1 | Farm | C | 1 | 1 | FC1 | NA | NA | 1 |
| 76-FD2-1 | Farm | D | 2 | 1 | FD2 | NA | NA | 2 |
| 77-A4 | Garden | NA | NA | NA | A4 | 4 | A | 2 |
| 78-FA1-1 | Farm | A | 1 | 1 | FA1 | NA | NA | 2 |
| 79-FC1-1 | Farm | C | 1 | 1 | FC1 | NA | NA | 2 |
| 81-S1 | Garden | NA | NA | NA | S1 | 1 | S | 2 |
| 82-FA3-2 | Farm | A | 3 | 2 | FA3 | NA | NA | 2 |
| 83-FC1-2 | Farm | C | 1 | 2 | FC1 | NA | NA | 1 |
| 84-FD2-2 | Farm | D | 2 | 2 | FD2 | NA | NA | 2 |
| 85-A5 | Garden | NA | NA | NA | A5 | 5 | A | 2 |
| 86-FA1-2 | Farm | A | 1 | 2 | FA1 | NA | NA | 2 |
| 87-FC1-2 | Farm | C | 1 | 2 | FC1 | NA | NA | 2 |
| 88-S2 | Garden | NA | NA | NA | S2 | 2 | S | 2 |
| 89-FA3-3 | Farm | A | 3 | 3 | FA3 | NA | NA | 1 |
| 90-FC1-3 | Farm | C | 1 | 3 | FC1 | NA | NA | 1 |
| 91-FD2-3 | Farm | D | 2 | 3 | FD2 | NA | NA | 1 |
| 92-W1 | Garden | NA | NA | NA | W1 | 1 | W | 2 |
| 93-FA2-1 | Farm | A | 2 | 1 | FA2 | NA | NA | 2 |
| 94-FC2-1 | Farm | C | 2 | 1 | FC2 | NA | NA | 2 |

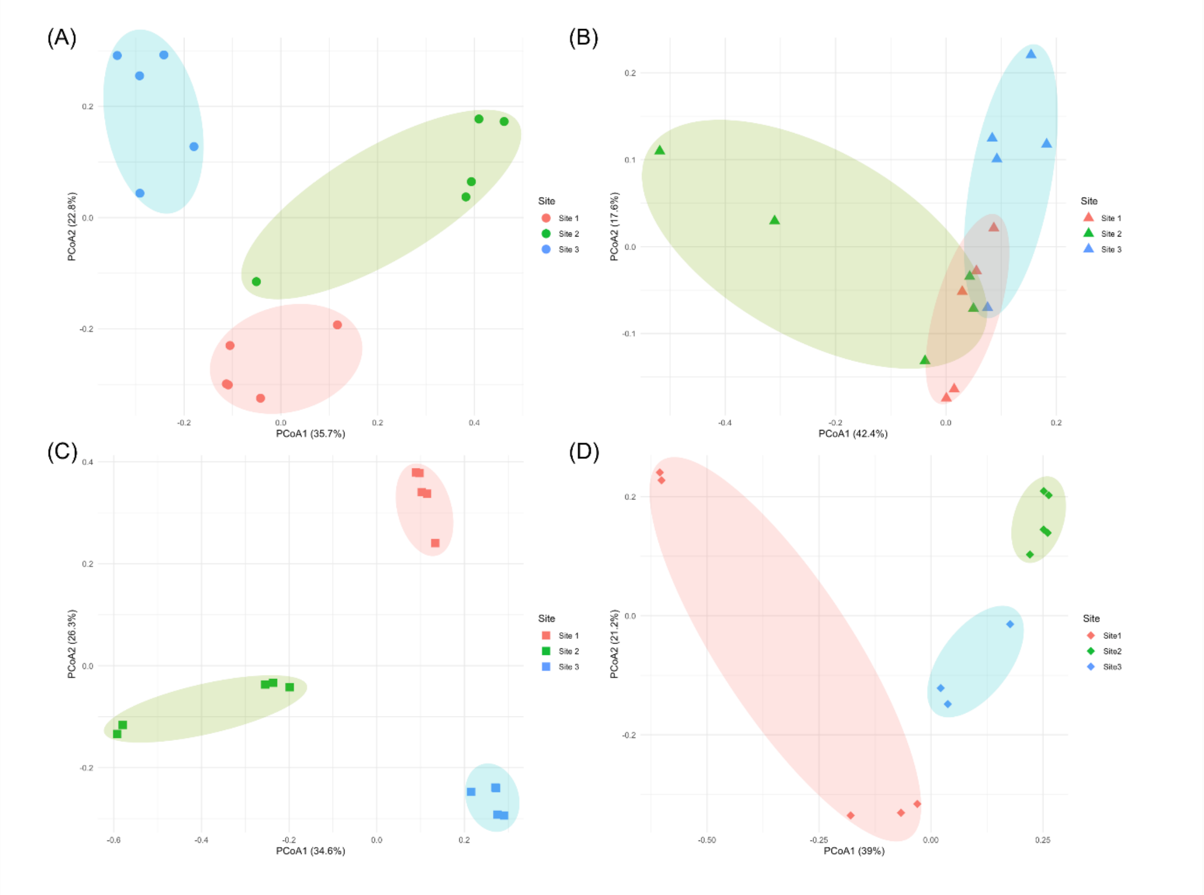


**Figure S4 PCoA Plots of Fungal Community Composition Across Different Sites Within Each Farm** Each plot (A-D) corresponds to a different farm, with samples grouped by site (Site 1-3). Colours are consistent across plots to represent site numbers but do not reflect comparable environmental conditions between farms. For this reason, comparisons should be interpreted only within individual farms, not across farms.

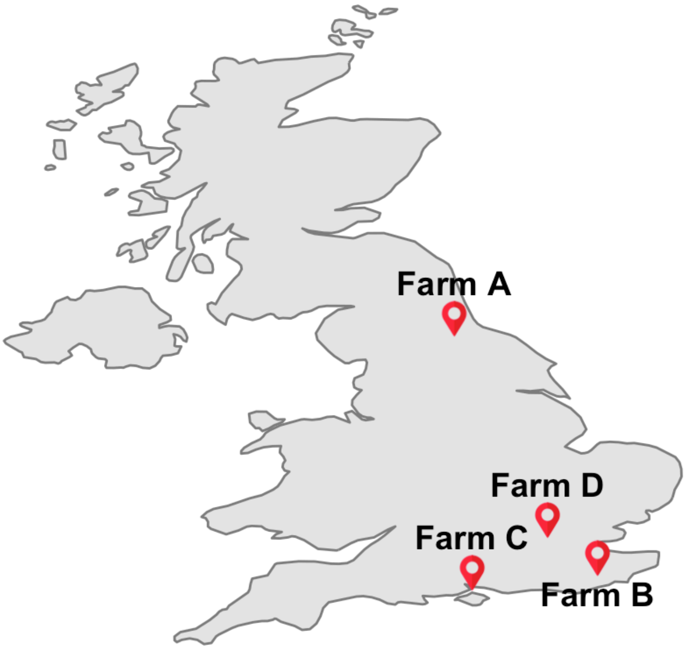


**Figure S5 Geographic Locations of Farm Sampling Sites in the UK** The four farms (A–D) represent spatially distinct sites used to investigate airborne fungal community variation.

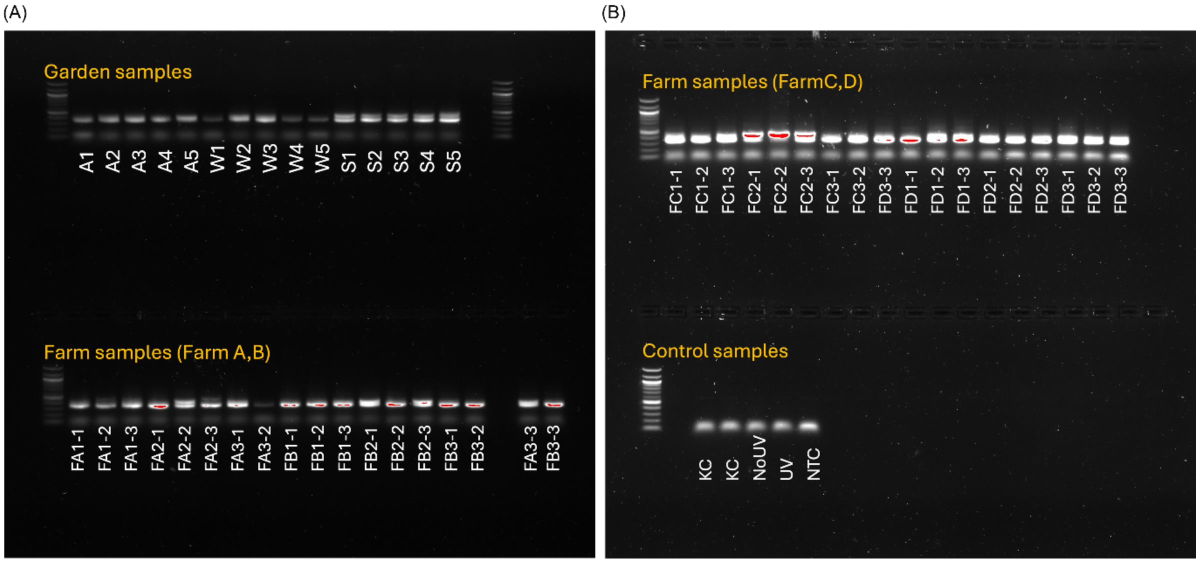


**Figure S6 PCR Verification of Environmental Samples Prior to Sequencing** Gel electrophoresis results showing successful amplification of ITS2 regions from garden samples, farm samples, and control samples. Amplicon presence was confirmed prior to sequencing. No amplification was observed in any of the controls (UV, NoUV, and NTC), indicating the absence of contamination. Garden samples include seasonal sets (A = autumn, W = winter, S = spring). Farm samples are labelled according to farm ID, site, and subsample number. Detailed sample ID conventions and metadata are provided in Supplementary Table S8.
